# Narratives for positive nature futures in Europe

**DOI:** 10.1101/2024.10.04.616255

**Authors:** Alessandra D’Alessio, Claudia Fornarini, Nestor Fernandez, Anandi Sarita Namasivayam, Piero Visconti, Jeremy Dertien, Maria Hällfors, Martin Jung, Francisco Moreira, Louise O’Connor, Matea Osti, Laura C. Quintero-Uribe, Martina Marei Viti, Henrique M. Pereira, Peter H. Verburg, Carlo Rondinini

## Abstract

1. The Nature Futures Framework (NFF) is a novel tool for the development of positive scenarios centred on the relationship of nature and people, emphasising biodiversity as part of the solution to environmental challenges across various spatial and temporal scales, explicitly addressing a plurality of values for nature.
2. In this work, we describe the process that has led to the formulation of continental-scale positive narratives for conservation in Europe based on the NFF and its value perspectives (Nature for Nature; Nature for Society; Nature as Culture), through a stakeholder group elicitation. We focused on 6 topics in the narratives: Nature Protection and Restoration; Forestry; Freshwater Ecosystems; Urban Systems; Agriculture, and Energy. We analyse differences and similarities among the narratives across these topics.
3. We develop three novel Nature Futures narratives for Europe with contrasting perspectives and priorities for the six topics. Within the EU socioeconomic trends and policy framework, common solutions that simultaneously tackle biodiversity conservation and instrumental and cultural Nature’s Contributions to People (NCP) provision emerged.
4. This set of narratives may integrate preferences concerning EU-level conservation targets and plausible socio-ecological development pathways, supporting the modelling of positive scenarios for nature that can be crucial in guiding policy decisions towards recovery of nature.

## 1. Introduction

The global biodiversity crisis has received increasing attention globally, but the actions have so far been insufficient to reverse the trend of declining biodiversity (CBD secretariat, 2020; IPBES, 2019). In Europe, the EU Biodiversity Strategy for 2030 provides a framework for current and future conservation endeavours by setting clear targets and objectives that largely align with the Kunming-Montreal Global Biodiversity Framework (EC, 2020a; KM GBF, 2022). The strategy sets ambitious goals, including the expansion of protected areas (PAs) to reach a minimum of 30% spatial coverage for both land and sea. Importantly, at least one third of these areas should be managed under strict protection. In addition, the pending European Nature Restoration Law demands action to ecologically restore at least 20% of degraded land and sea areas within the EU, and support the recovery of ecosystems and species in synergy with area protection targets (EC, 2022a). Yet, the long history of intensive exploitation of ecosystems in Europe and conflicts with other relevant socio-economic activities, such as agricultural, forestry, urbanisation or energy production, makes the achievement of these policy targets challenging.

Achieving ambitious goals in the context of competing interests requires an integrated management approach that explores all relevant nature conservation values and options. Environmental change scenarios are valuable for nature conservation for investigating the potential impacts of different societal development pathways and policy choices on biodiversity and Nature’s Contributions to People (NCP), while also facilitating communication and involving multiple stakeholders in the process (Pereira et al., 2020). The widely used Shared Socio-Economic Pathways (SSPs) scenario framework integrates drivers such as demography, governance efficiency, inequality at both national and international levels, socio-economic advancements, institutional factors, technological advancements, and environmental conditions (van Vureen et al., 2014; O’Neill et al., 2014). However, scenarios based on SSPs typically do not take in consideration positive features specifically for nature and biodiversity, and are thus limited in their use for exploring different societal preferences related to the role of nature, and related policies driving human socio-economic development (IPBES, 2016; Saito et al., 2019; Pereira et al., 2020; Lundquist et al., 2021).

At the same time, it is increasingly clear that different stakeholders exhibit different preferences for nature, depending both on their relationship with nature and the information provided given different nature management options (Capper et al., 2024; Carvalho Ribeiro et al., 2013; van der Wal et al., 2014). Recognizing the plurality of views of nature across people is important to democratise the management of landscapes, acknowledging tensions between stakeholders but also their perspectives on nature (Dotson & Pereira, 2022). This richness of perspectives on nature is not currently represented in existing scenarios, with often only one “desirable” perspective for nature being considered in a given set of scenarios (Rosa et al., 2017; Pereira et al., 2020).

To address the limitations within existing scenarios, the expert group on scenarios and models of the Intergovernmental Science-Policy Platform on Biodiversity and Ecosystem Services (IPBES) developed the Nature Futures scenario Framework (NFF) (IPBES, 2023a). The NFF aims to support the development of positive scenarios centred on the relationship of people with nature across various spatial and temporal scales (IPBES, 2023b; Kim et al., 2023). This framework incorporates different perspectives, all with nature at the centre of the scenario design rather than just as an outcome, and allows the consideration of diverse value perspectives (Rosa et al., 2017; Pereira et al., 2020). NFF scenarios encompass three value perspectives that capture and cluster the many different preferences for nature across people (Mansur et al., 2022; Pascual et al., 2023), and can be represented as three corners of a triangle (Fig. S1). The Nature for Nature (NfN) perspective, emphasises the intrinsic value of nature, including preserving individual species and species diversity, habitats, ecosystems, natural processes, and the self-regulatory processes of nature. The Nature for Society (NfS) perspective focuses on the maximisation of instrumental values, benefits, and services that biodiversity and ecosystems provide to people, including food provisioning, water purification, disease control. Finally, the Nature as Culture (NaC) perspective highlights the relational values between nature and people, where society, traditions, beliefs and emotions drive socio-ecological landscapes, such as silvo-pastoral landscapes (Bugalho et al., 2011; Zerbe, 2022).

The NFF has been applied to assess preferences for nature in existing participatory scenarios (Quintero-Uribe et al., 2022), to develop new scenarios, e.g., in a National Park in the Netherlands (Kuiper et al., 2022), in a rural landscape in northeastern Japan (Haga et al., 2023), and in urban management (Mansur et al., 2022). Recently, the framework has been adopted to explore how contrasting narratives would translate into land use scenarios for Europe by 2050 (Dou et al., 2023). However, the NFF has never been applied to formulate continental-scale positive nature future narratives. These aim to integrate societal visions and preferences concerning EU-level conservation targets and plausible socio-ecological development pathways, thus supporting policy decisions towards recovery of nature.

Here we designed NFF narratives for Europe through a participatory approach with stakeholders that were previously identified through a mapping exercise, and then invited to join two stakeholder engagement events, both in person and online. The narratives describe different scenarios that explore conservation and restoration priorities and policies. We aimed to answer the questions: what are possible contrasting positive futures for European landscapes? What are the common enabling conditions that need to be met for any of these positive futures to come to fruition? Through a participatory process, we gathered perspectives and priorities from stakeholders and formulated NFF narratives based on key topics: Nature Protection and Restoration, Forestry, Freshwater Ecosystems, Urban Systems, Agriculture, and Energy. These topics emerged in the context of the current challenges for nature conservation to help envision a sustainable future for nature and society. The narratives can support integrated planning and land use modelling towards the achievement of EU policy targets, by supporting modellers in the field of conservation, and consequently assisting the EU Member States in developing an ecologically representative, resilient, and well-connected Trans-European Nature Network (TEN-N) (NaturaConnect, 2024). To our knowledge they are the first of their kinds that explicitly place conservation and restoration in the centre, in line with EU policy targets and in a globally comparable framework (IPBES NFF).

## 2. Material and methods

To develop the NFF narratives aligned with the three perspectives, representing the corners of the triangle (Fig. S1), we implemented the method from Pereira et al. (2020), into a sequence of ten steps (Fig. 1) (see Appendix 2 for further details).

**Figure 1.**
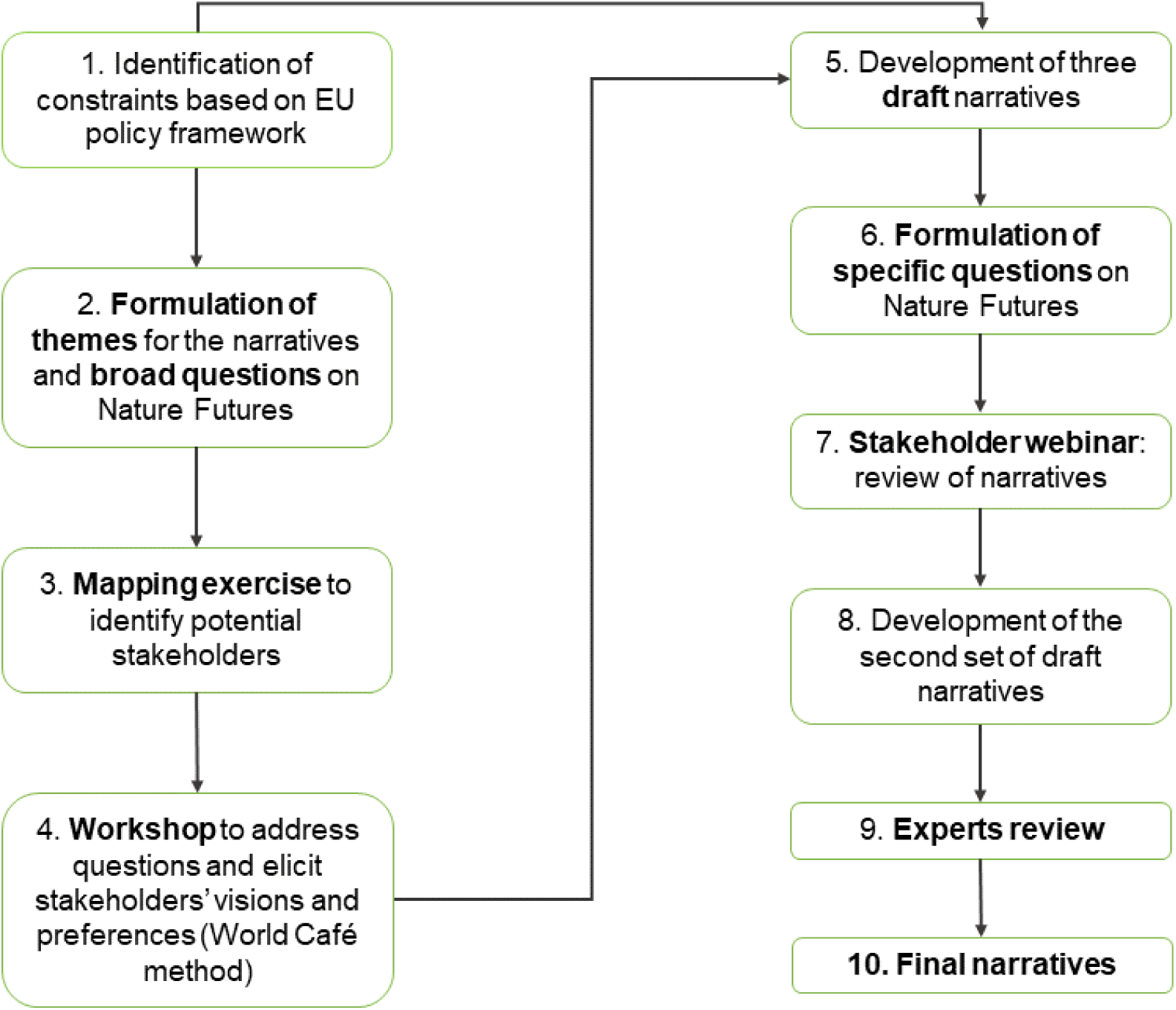
The process of development of the Nature Futures narratives for Europe.

**1)** We identified a set of EU assumptions, or ‘constraints’, that coerce the NFF narratives. We considered key EU legislation, regulations, objectives and strategic priorities as mandatory for all NFF narratives. These include the EU Biodiversity Strategy objectives for 2030, such as the expansion of PAs and strictly protecting one third of these areas; the implementation of multifunctional Green and Blue Infrastructure; and the Nature Restoration Law (EC, 2022a). We also took into account the Common Agricultural Policy; the EU Farm to Fork Strategy (EC, 2020b); the “No Net Land Take ‘’ by 2050 objective (EC, 2016); and the European Climate Law (EC, 2023b). **2)** According to the challenges and constraints facing Europe, we decided to address a preliminary set of themes and, based on them, we formulate a set of broad questions to be asked to stakeholders (Appendix 2.1). 3) We identified key stakeholders through a mapping exercise, based on their influence in specific sectors of interest at the European level. Stakeholders were mapped using a power-interest grid, a conceptual framework that sorts stakeholders into four quadrants based on their interest in the different workflows of the process and influence on its outcomes (Figure S2; Appendix 2.2)

This categorization helps determine the key stakeholders, with high power and high interest, that should be deeply involved in stakeholders elicitation processes to identify plausible and supported Nature Future narratives that are compatible with the achievement of the objectives of the EU Biodiversity Strategy 2030.

**4)** In a second phase, we organised an in-person workshop with stakeholders to elicit their perspectives on the future of nature. We held a three-day in-person workshop (Leipzig, Germany, 8-10 May 2023). During this event, scientists of the NaturaConnect project were introduced as internal stakeholders, representing several expertise within the conservation sector. The workshop aimed to gather insights on the future of nature in Europe, using the World Café method for structured dialogues led by moderators (Brown, 2010) (Appendix 2.3).

The first World Café round, which focused on landscape changes, agriculture management, and conservation motivations, was facilitated by showing pictures of different European landscapes, selected according to the themes identified in the previous step. Participants moved between tables that represented the different corners of the NFF triangle to envision future European landscapes contrasting the three different NFF perspectives on nature. Subsequently, the discussion moved into the previously defined themes (Appendix 2.3). **5)** This visioning exercise was propaedeutic to develop the first draft of the narratives, by elaborating and revising the outcomes with moderators of each workshop’ session. After the workshop, indeed, we refined the three narratives “Nature for Nature”, “ Nature for Society” and “Nature as Culture”, focusing them on six main recurring topics: Urban systems, Forestry, Freshwater Ecosystems, Energy, Agriculture, and Nature Protection and Restoration (Appendix 2.3). **6)** Since gaps concerning preferences and different perspectives emerged, particularly on Nature Protection and Restoration topics, we defined additional questions on nature futures to improve the narratives (Appendix 2.4). **7)** A draft version of the narratives was presented during a 2-hour public webinar (4 July 2023). It served to harvest additional feedback and insights, through 15 interactive questions via Mentimeter (www.mentimeter.com), following each narrative presentation (Appendix 2.5). **8)** After the webinar, the most frequent remarks and new information were collected. Thus, both stakeholders’ event inputs were analysed and integrated to create a coherent second set of draft narratives. **9)** Finally, following a further review by the experts group of the NaturaConnect project, **10)** we developed a final set of narratives (Appendix 2.5). The study has been approved by the NaturaConnect committee which has ensured the ethical requirements and that all people involved in the stakeholders’ event gave their informed consent for participation and to share the obtained outcomes.

We analysed the main differences and commonalities across the narratives and we highlighted contrasts across the narratives concerning the six topics. Specifically, we analysed some specific aspects involving the six topics that were key in distinguishing the NFF narratives: the dichotomy between land-sharing and land-sparing, the restoration approach, the importance of maintaining the integrity of freshwater resources, the level of forest management, the human presence in protected areas, the population flow and the urban configuration, the agricultural strategies and the implementation of wind and solar energy. Reflecting the importance of these aspects in each narrative, we attributed each a gradient of preference from Minimum to Medium to Maximum.

## 3. Results

The in-person stakeholder workshop was joined by 41 participants from 13 European countries, including 13 external stakeholders and 28 conservation scientists and practitioners from the NaturaConnect project. All participants represented institutions and stakeholder groups of the European environment conservation (95,4%) hunting (2,3%) and land use planning (2,3%) sectors.

The webinar brought together a group of 115 participants from 18 countries, all European except one. The stakeholders who responded to the specific question (68 people) gave 100 answers, about the sector they belong to. This means that some people are declared to belong to more than one sector. The sectors are distributed as follows: nature conservation (54%), land use planning, (13%), forestry (9%), social science (8%), policy and law (5%), urban (3%), marine (2%), agriculture (1%), tourism (1%) and other sectors (4%). Based on the webinar participants’ responses (60%), 80% belonged to nature conservation governmental or non-governmental organisations. However, it should be noted that 35% of participants who participated in the webinar their affiliated entity and sector remained unknown.

Through the stakeholder elicitation and refinement by the expert group, we designed three narratives that describe different nature futures in Europe, one per each corner of the NFF triangle: Nature for Nature (Box 1), Nature for Society (Box 2) and Nature as Culture (Box 3). Below we summarised the main content of each narrative by topic (Table 2) and we highlighted the differences and commonalities among the narratives.

**Table 1:**
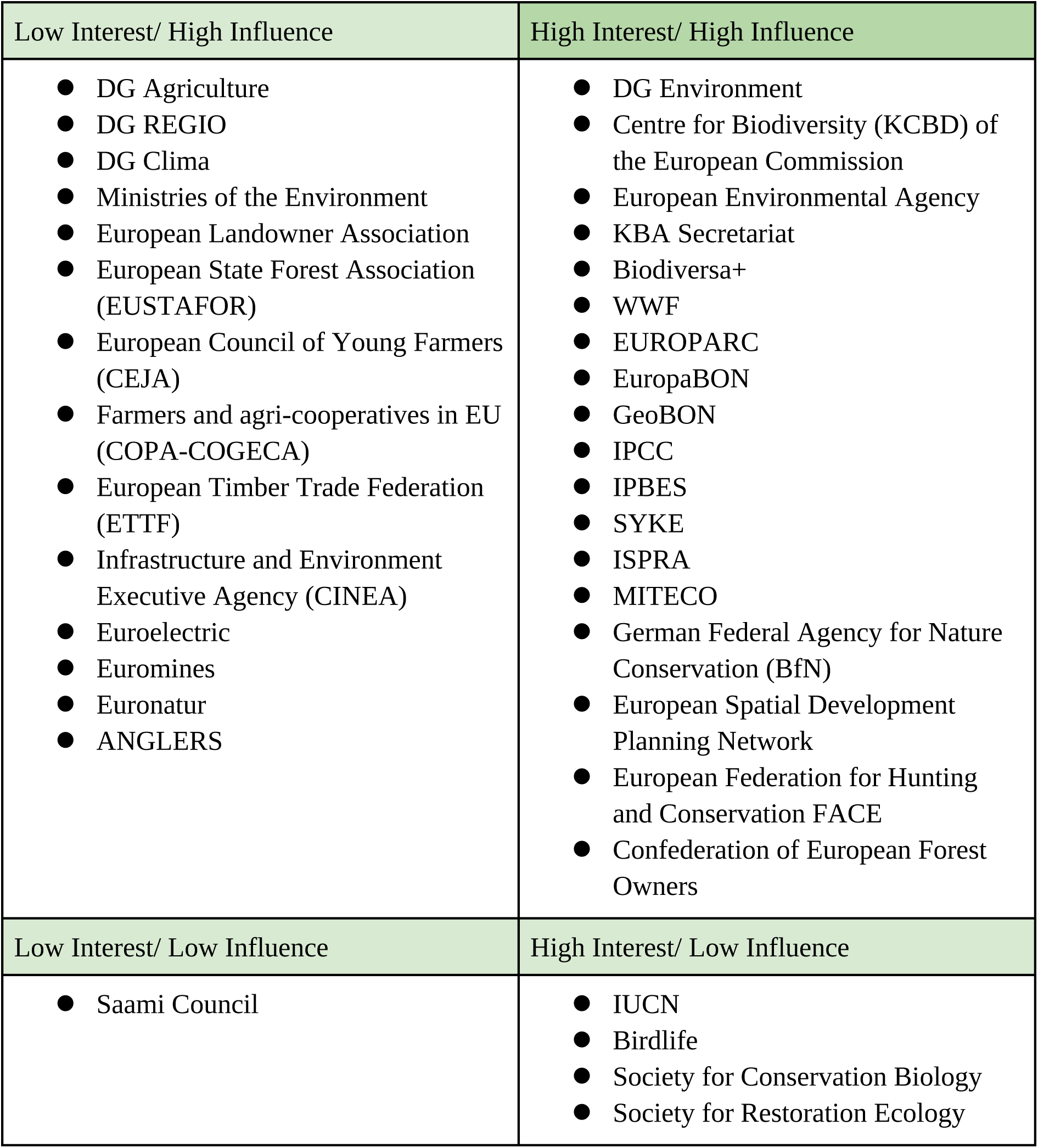
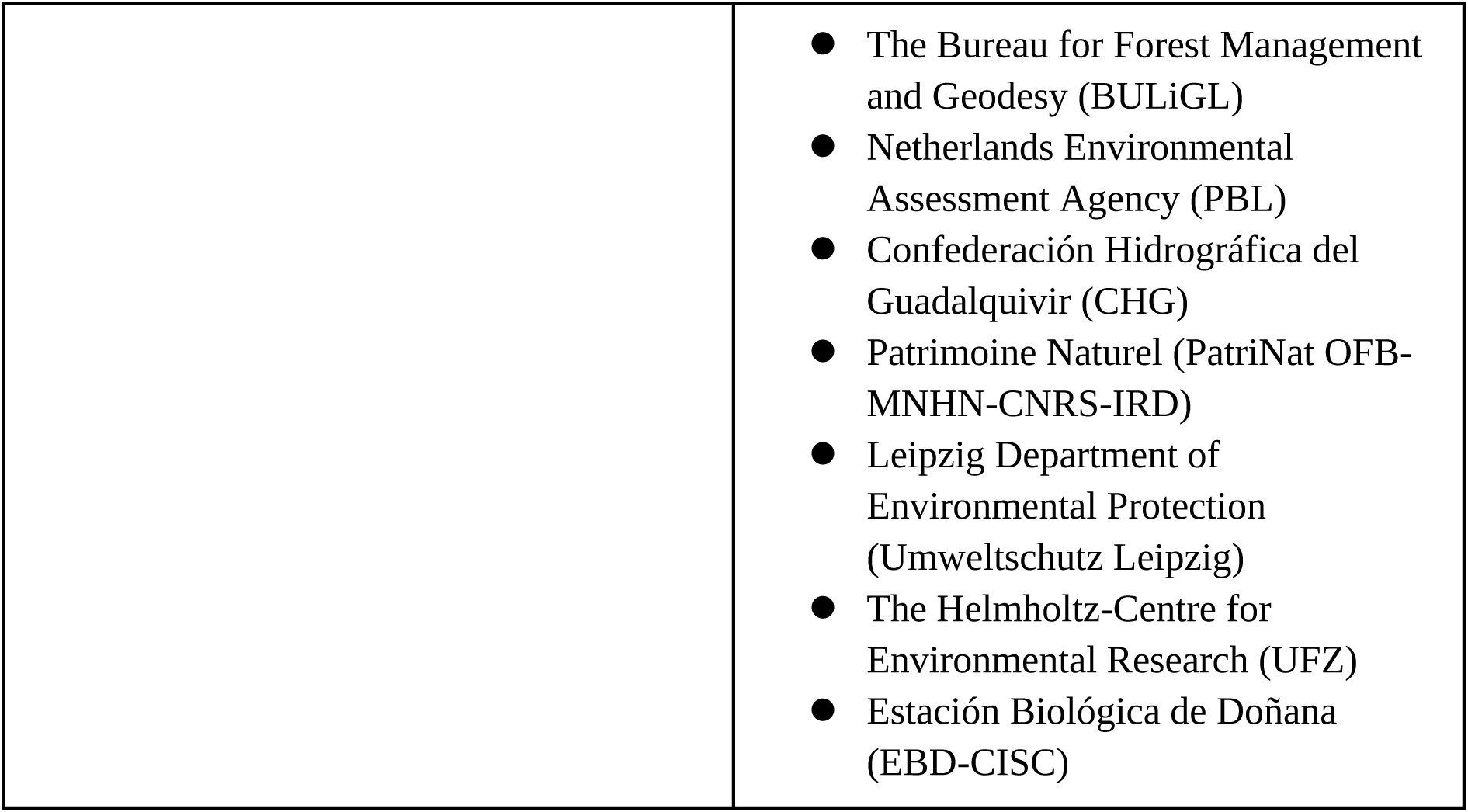
Table listing the stakeholders identified through the mapping exercise and clustered according to their levels of interest and influence. Key stakeholders, with high interest and influence are reported in the top right-hand box.

| Low Interest/ High Influence | High Interest/ High Influence |
| --- | --- |
| ● DG Agriculture<br>● DG REGIO<br>● DG Clima<br>● Ministries of the Environment<br>● European Landowner Association<br>● European State Forest Association (EUSTAFOR)<br>● European Council of Young Farmers (CEJA)<br>● Farmers and agri-cooperatives in EU (COPA-COGECA)<br>● European Timber Trade Federation (ETTF)<br>● Infrastructure and Environment Executive Agency (CINEA)<br>● Euroelectric<br>● Euromines<br>● Euronatur<br>● ANGLERS | ● DG Environment<br>● Centre for Biodiversity (KCBD) of the European Commission<br>● European Environmental Agency<br>● KBA Secretariat<br>● Biodiversa+<br>● WWF<br>● EUROPARC<br>● EuropaBON<br>● GeoBON<br>● IPCC<br>● IPBES<br>● SYKE<br>● ISPRA<br>● MITECO<br>● German Federal Agency for Nature Conservation (BfN)<br>● European Spatial Development Planning Network<br>● European Federation for Hunting and Conservation FACE<br>● Confederation of European Forest Owners |
| Low Interest/ Low Influence | High Interest/ Low Influence |
| ● Saami Council | ● IUCN<br>● Birdlife<br>● Society for Conservation Biology<br>● Society for Restoration Ecology |
|  | ● The Bureau for Forest Management and Geodesy (BULiGL)<br>● Netherlands Environmental Assessment Agency (PBL)<br>● Confederación Hidrográfica del Guadalquivir (CHG)<br>● Patrimoine Naturel (PatriNat OFB-MNHN-CNRS-IRD)<br>● Leipzig Department of Environmental Protection (Umweltschutz Leipzig)<br>● The Helmholtz-Centre for Environmental Research (UFZ)<br>● Estación Biológica de Doñana (EBD-CISC) |

**Table 2:** Summary of the narratives. The main content of the narratives (in column) is described per topic (in rows). Note that the topic Nature Protection and Restoration includes a focus on conservation goals for each narrative, describing the Protected Areas (PAs) aim and use, and the restoration strategy. Restoration is also the main focus in the Freshwater Ecosystem row. Forestry and agriculture topics are focused on different land management approaches, while Urban Systems and Energy address infrastructures development that also involve people distribution.

| Topic | Nature for Nature | Nature for Society | Nature as Culture |
| --- | --- | --- | --- |
| Nature Protection and Restoration | Emphasis on ecological integrity and resilience. Irreplaceable and particularly vulnerable species and ecosystems receive high priority.<br>In protected areas (PAs), activities are minimised in line with biodiversity conservation objectives. In strictly protected areas no management and no intervention is carried out in sites with high ecological integrity. | Emphasis on Nature’s Contributions to People (NCP) provisioning and associated species and ecosystems.<br>In PAs, there is moderate to high tolerance for human activities/ intervention related to Nature’s Contributions to People use. In strictly protected areas focus is on preserving ecosystems for which the processes and functions associated with NCP depend on minimal disturbance. | Emphasis on cultural landscapes, including high nature value farmland and associated species.<br>In PAs, there is high tolerance for cultural human activities. In strictly protected areas focus is on culturally relevant species and ecosystems which require minimum disturbance. |
|  | Passive restoration is enhanced. Structural and functional connectivity is improved for all species through Green and Blue Infrastructures. | Active restoration is enhanced. Ecosystems’ connectivity that supports NCP provision is improved, especially in peri-urban landscapes and across cultivated land through Green and Blue Infrastructures. | Active restoration is enhanced. Connectivity is improved for symbolic species and cultural landscapes, especially agroecological areas with hedgerows and natural patches, and cities through Green and Blue Infrastructures. |
| Forestry | Land sparing approach with no logging in old-growth forests. Passive afforestation through natural succession enhances the complexity of forests. | Land sharing approach with forests managed to have multifunctionality, maximising NCP and biodiversity. Active afforestation with native species that provide NCP. | Land sharing approach with local communities managing forests to provide cultural services. |
| Freshwater Ecosystems | Restore freshwater ecosystems maximising ecological integrity by removing obsolete barriers for species connectivity and ecological flows. | Restore freshwater ecosystems that provide NCP and minimise barriers' impacts on biodiversity and NCP, such as flood mitigation. | Restore freshwater ecosystems with cultural/traditional value or areas linked to emblematic species. |
| Agriculture | Land sparing approach to save more space for biodiversity conservation. | Land sharing/sparing mixed approach. Large-scale farming and NBS to provide NCP. | Land sharing approach to better integrate nature with anthropogenic traditional activities of cultural value. |
| Urban Systems | High-rise compact cities but no sprawl with population flow from rural areas to cities. | Moderately compact cities that maximise access to NCP. | No high-rise compact cities and increased population flow from cities to rural areas. |
| Energy | Renewable Energy implementation avoids areas of conservation concern. | Renewable Energy Sources are planned to reduce land-take impacts on biodiversity and related NCP. | Renewable Energy plants are placed in isolated areas to avoid culturally important places and landscapes. |

### 3.1 Differences among the narratives

The main difference among the narratives is the preference towards a land sharing or sparing approach, across several topics such as Agriculture, Urban System, Forestry and Energy (Fig. 2).

**Figure 2.**
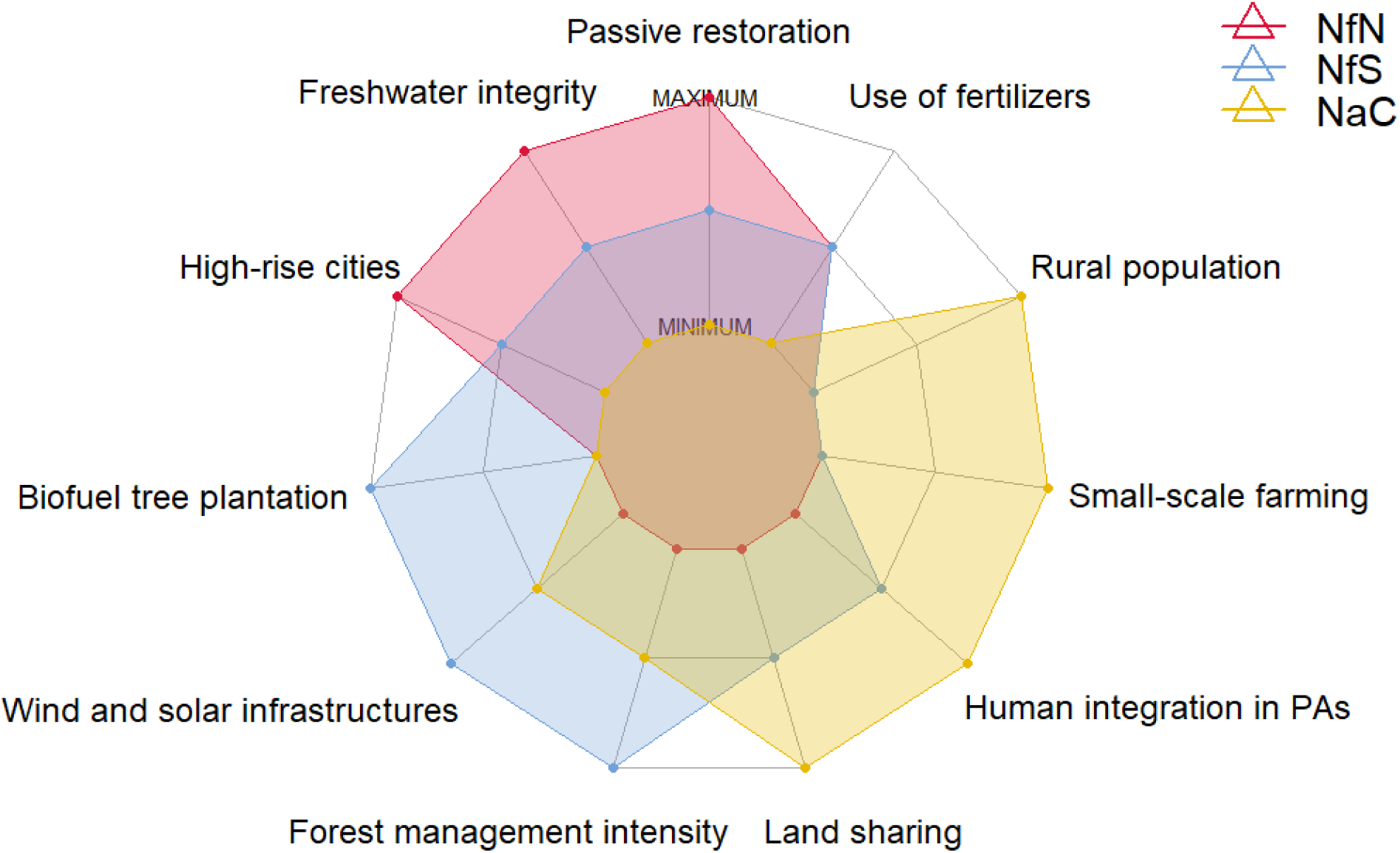
Spider diagram showing the main differences among the Nature Futures for Europe. The red, blue, and yellow polygons represent NfN, NfS and NaC, respectively. Axes represent a gradient measured on an ordinal scale from Minimum to Medium to Maximum. This gradient reflects stakeholders preferences for all NFF corners, on topics selected for drafting the narratives (Nature Protection and Restoration, Agriculture, Urban Systems, Freshwater Ecosystems, Forestry, Energy).

In the NaC perspective, land sharing is preferred (Box 3), whereas, in NfN, land sparing is favoured (Box 1). NfS requires a moderate gradient of land sharing to provide NCP (Box 2). Therefore, large-scale agriculture is practised in both NfS and NfN, and small-scale farming resulted as preferred in NaC. Stakeholders found elements of ecological integrity as cross-cutting elements in NFF, whereas in the other narratives it is not perceived as significant. In this context, freshwater ecosystems protection and restoration seem to be crucial within the NfN narrative, while being less considered in NfS and NaC where they reach the lowest value. In the NfN perspective, human activities are minimal in PAs because access to these areas is limited, but are expected to be moderate in NfS and maximal in NaC, where they are located near human settlements to improve accessibility (Fig. 2). In NfN, passive restoration is preferred and forests are less managed than in NaC and NfS. Development of high-rise compact cities is at its maximum in NfN to make space for nature. A similar urban development occurs in NfS. Conversely, in NaC, people move from large cities and peri-urban areas to medium and small settlements in rural areas with low population density. In NfN, ecological integrity and connectivity have priority over renewable energy sources such as wind and solar farms. In contrast, nature has low priority over renewable energy sources implementation in NfS, while being moderate in NaC. Fast-growing tree plantations for biofuel production (e.g. poplars) are more encouraged in NfS than in the other narratives (Box 2). The amount of space required for this activity results in no forest patches allotted for biofuel in NfN and NaC (Box 1) (Box 3).

### 3.2 Commonalities

Some common concepts emerged across the narratives, since they were all based on the 2030 EU Biodiversity goals, and included mutually beneficial solutions that address biodiversity conservation and NCP provisioning (Fig. 3).

**Figure 3:**
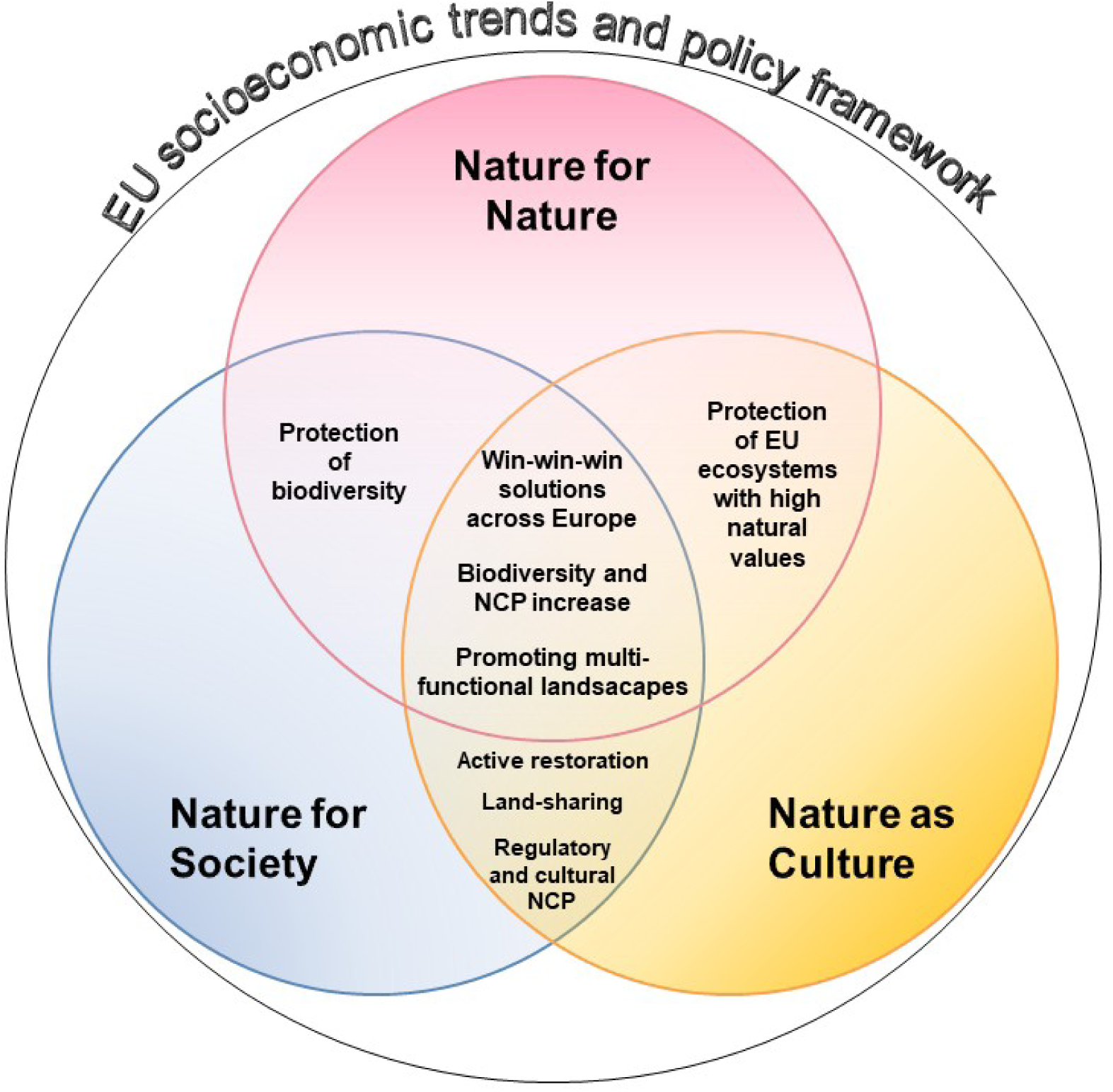
Venn diagram showing the commonalities (coloured in black) among the Nature Futures for Europe. Overall, win-win-win solutions and an increase in biodiversity and Nature’s Contributions to People (NCP) are envisioned for all NFF corners.

Restoration efforts can achieve multiple objectives for nature and people by enhancing ecosystem integrity and connectivity, and simultaneously ensuring the practical uses and cultural values of nature. For example, restored natural areas along rivers may provide umbrella habitats and regulate flooding whilst also creating space for recreational activities (Fig. 3). Infrastructure planning, including highways, railways, and renewable energy plants, aims to improve coexistence between humans and nature for space efficiency, though minimising impacts on species and ecosystems. Energy communities, which are organisations that rely on sharing energy among local citizens, public administrations, and enterprises (EC, 2023a), may reduce the need for linear energy infrastructures. The deployment of photovoltaic panels on roofs could allow saving space outside urban areas. Urban greening and gardening initiatives may reduce the human carbon-footprint and ensure environmental sustainability, NCP, biodiversity and connectivity. The implementation of zero-emission public transportation and bike pathways within and around cities is a shared measure to mitigate climate change effects, contributing to the improvement of both nature and human health.

Promoting multifunctional landscapes is central in NfS and NaC, indeed sustainable management of agricultural and forest landscapes may support various functions concurrently, such as the optimisation of biofuel production through the use of crop and wood residues. Sustainable forestry is also beneficial in terms of carbon sequestration and availability of recreational areas, and it supports the maintenance of biodiversity, and its productivity, vitality, regenerative capacity, as well as the provisioning, over time, of material and regulatory NCP.

## Box 1. Nature for Nature (NfN)

In the NfN narrative, the value of nature is intrinsic and independent from any direct benefits that people may gain from nature. The protection and restoration of the ecological integrity of ecosystems are therefore key priorities in this narrative and thereby land sparing approaches are pursued. Strict protection is envisioned for natural areas to preserve the integrity and resilience of nature within the European protected area network. Thus, human activities are minimised in PAs as access to these areas is restricted. Conservation focuses on sensitive and irreplaceable species and habitats. Both structural and functional connectivity is improved for all species through Green and Blue Infrastructures. Restoring and ensuring the connectivity of PAs is a priority pursued to help recover the characteristic ecological flows of undisturbed ecosystems. Restoration of connectivity in freshwater ecosystems is essential in this narrative and obsolete dams are removed for this purpose. Natural forest dynamics is promoted, thus enhancing both structural and functional complexity and natural regeneration and turnover. Forest harvesting is reduced to a minimum, especially in old-growth forests and in strictly protected areas. To leave space for nature conservation, high-intensity agriculture is maintained to maximise production without expanding agricultural land. Precision farming is promoted to minimise impacts from agriculture. No increase in urban sprawl but high-rise compact cities development are deemed desirable. Renewable energy production, such as wind and solar, is established outside areas with high biodiversity values, also excluding buffer zones around PAs and other sensitive conservation areas. They are strategically placed in already degraded areas and high-intensity agricultural landscapes. Power lines are constructed along pre-existing infrastructures, and efforts are made to conceal them underground to minimise wildlife mortality and disturbances.

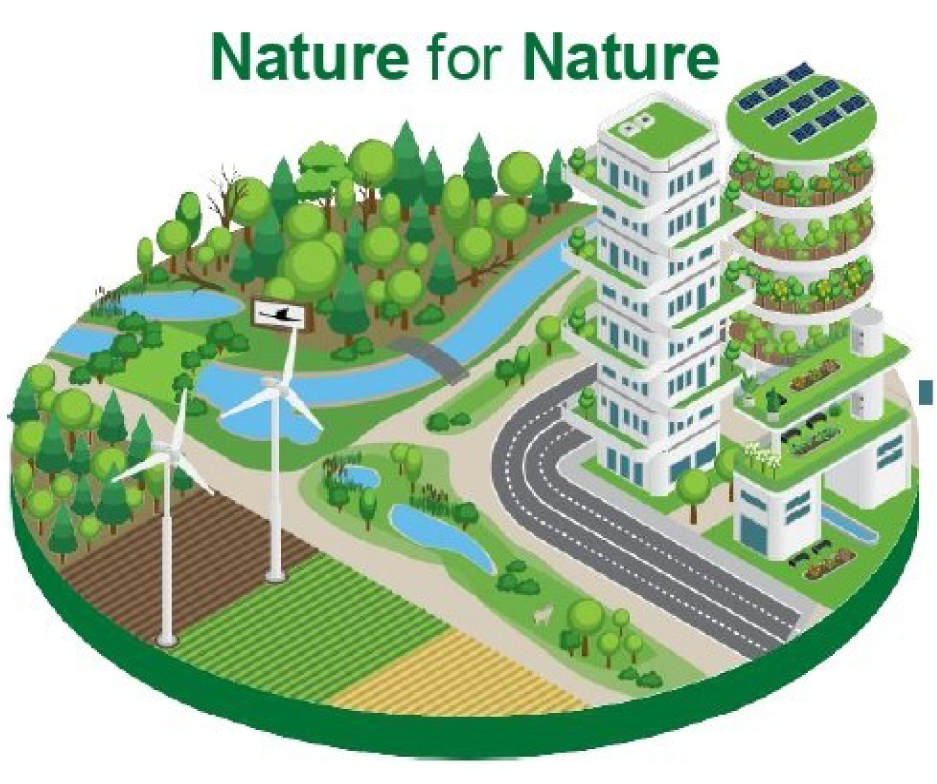

## Box 2. Nature for Society (NfS)

In the NfS perspective, emphasis is placed on the instrumental value provided to people. As a result, ecosystems are protected and restored with the aim of boosting the provisioning of NCP. To allow this provisioning, PAs are located where both NCP supply and demand are high and human activities are moderate. Species conservation is a priority mainly when it is associated with the supply of a specific NCP. Ecosystems for which the processes and functions associated with NCP depend on minimal disturbance are strictly protected. Ecological corridors are designed and restored taking into account their capacity to provide multiple benefits to people, especially in peri-urban landscapes and across cultivated land through Green and Blue Infrastructures (EC, 2019). Overall, active management and restoration approaches are used to prevent natural hazards (such as fire and flood risk) or reverse their impacts, promote carbon sequestration and sustainable timber extraction in forests, guarantee good water quality and supply, and ensure wild fish supply in freshwater ecosystems. Moderate land sharing is necessary for providing NCP in NfS. High-intensity agriculture and farming are away from areas of conservation concern. However, to enhance the co-benefits related to NCP, such as increasing biodiversity that leads to a better provision of resources or services for society and providing agroecological landscapes for species and habitats of high conservation interest (e.g. farmland birds and Dehesas), agriculture is slightly de-intensified and often integrated with NBS (e.g., hedges, green linear elements, restoration of landscape complexity). Moderately compacted urban areas are planned to facilitate beneficial contact between society and natural features, implying some urban sprawl in peri-urban areas. The provision of renewable energy is given priority over nature; thus, dams are managed to have minimal impacts on biodiversity and NCP (e.g., flood regulation, sediment retention, water quality and control of invasive species). Among renewable energy sources, fast-growing tree plantations for biofuel production (e.g. poplar) are encouraged, and wind and solar power plants are planned to minimise potential impacts on the provision of NCP.

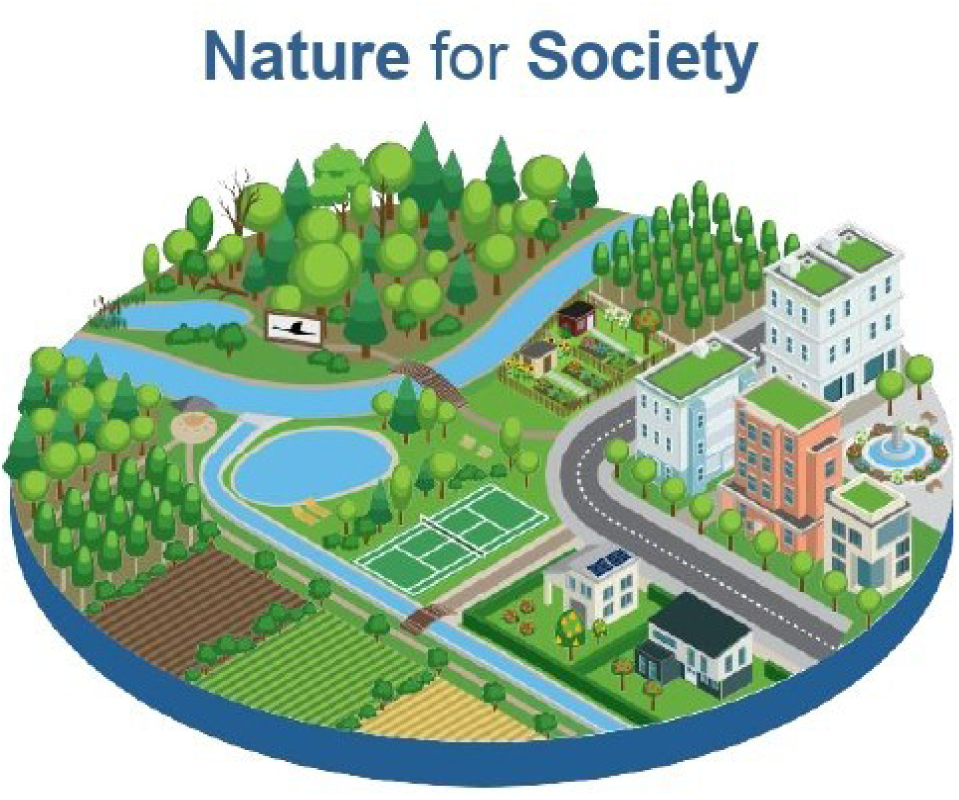

## Box 3. Nature as Culture (NaC)

The NaC narrative focuses on the relational values for nature, expressing personal and collective emotional connections that people have with nature. Therefore, human activities and presence within nature are tolerated more in this narrative than in the others. Strict protection focuses on culturally relevant species and ecosystems which require minimum disturbance. Overall, the protected area (PA) network is managed with a strong focus on maintaining culturally important practices, protecting heritage landscapes, and agroforestry and other human-modified systems with high natural value (Halada et al., 2011). These are done through initiatives such as UNESCO Man and Biosphere reserves (MAB) (Reed, 2019). Thus, traditional land use practices and experiences that connect people to specific landscapes are prioritised in PAs (e.g., Farm to Fork initiatives, wine routes, transhumance of livestock, high nature value farmland, biodiversity-friendly farming, pilgrimage routes, hiking and enjoyment of nature). Conservation efforts address species and habitats associated with culturally important activities, such as fishing or hunting, and the expansion of PAs aims to meet conservation objectives that preserve culturally valued species (e.g., migratory birds and fish, charismatic species), habitats (e.g. agroforestry systems, hay meadows), and ecosystem services. Landscapes of cultural, educational and/or historical importance and habitats of culturally important species are restored, and their connectivity is improved, with an additional aim to bring nature back to highly degraded areas, cities and agroecological areas through Green and Blue Infrastructures. Forests are managed by prioritising tree species with high cultural value. Ancient trees and other natural monuments are preserved. Freshwater ecosystems with a historical and cultural role, or those that are important for emblematic species, are also protected and restored, removing obsolete dams unless they have cultural importance. In agriculture, priority is given to the revitalisation of extensive and traditional agricultural practices in rural areas with high conservation and cultural value. These activities enhance the connection between nature and people that prefer living in rural areas, supporting the revitalization of small villages and regional towns. Renewable energy infrastructure is concealed from humans in order to preserve the aesthetics of the landscape.

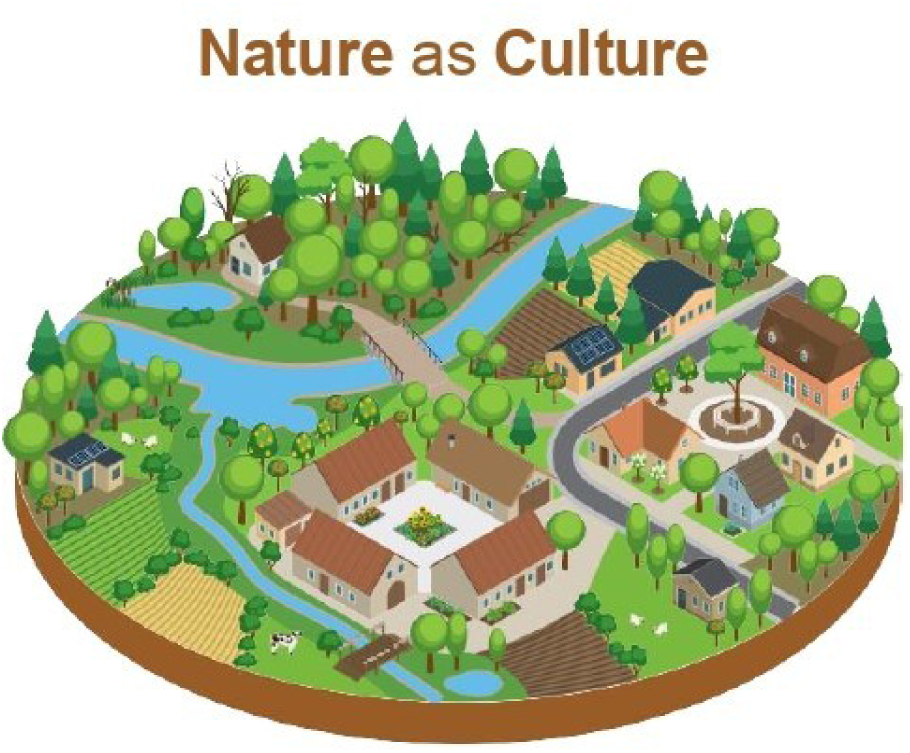

## 5. Discussion

Here, we formulated three NFF narratives, through a co-design approach carried out by scientists with a range of expertise who have elicited stakeholders’ preferences. This allowed us to produce scenarios that explore conservation and restoration priorities for achieving the European biodiversity targets for 2030, and can be applied for modelling positive futures for nature.

Our narratives highlight differences stemming from the three different sets of nature values that the NFF describes. The gradient of land sharing vs land sparing (Kremen, 2015) was the main axis for teasing the three NFF perspectives apart. This was evident across several topics such as Agriculture, Urban System, Forestry, and Energy. The stakeholders’ preferences were oriented toward land sharing in NaC (Box 3), based on the general expectation that land sharing that integrates people with nature can be beneficial in terms of recreation activities, carbon sequestration, pollination, livelihood, and biodiversity. The land-sparing approach is mainly useful to maintain the space allocated for spared reserves (Kremen, 2015) as emerged in the NfN narrative, focused on strict nature conservation (Box 1). Land sharing cannot achieve the conservation of all species, especially those more sensitive to human disturbance, and it has often been associated with lower species richness compared with land sparing (Edwards et al., 2013; Cannon et al., 2019; Balmford, 2021). Rural abandonment envisioned in the European NfN and NfS perspectives may lead to an increase in biodiversity, especially due to the abandonment of previously intensively managed land (Daskalova & Kamp, 2023). However, the opposite trend is already happening in some European countries: regions of Central and Eastern Europe are experiencing large human population flows from urban to rural areas (Toader et al., 2018; Despotovic et al., 2020). Nevertheless, land sparing may be difficult to achieve in most of the European context, as there is little land available to be fully ‘spared’ in the first place. In conclusion, the combination of context-specific land sharing and land sparing measures could be preferential when aiming to enhance biodiversity (Grass et al., 2021; Sidemo-Holm et al., 2021), and could be the best compromise to achieve sustainable targets for Europe.

Despite the differences, some common concepts emerged across the narratives based on the 2030 EU Biodiversity goals and targets, including mutually beneficial solutions for biodiversity and NCP (IPBES, 2016). Restoration efforts that enhance ecosystem integrity improve utilitarian functions such as water and air purification, pollination, climate change mitigation, and flood prevention, as well as the preservation of cultural values (Schindler et al., 2014; Zerbe, 2022). We considered multifunctional landscapes crucial in the NfS and NaC narratives (Fig. 4). Their importance recur in different sectors, such as agriculture and forestry (Renting et al., 2009; Lindroth et al., 2012; Diez & Garcia, 2012), as it has been pointed out across the NFF perspectives.

Efficient and carefully planned infrastructures, including renewable energy production and urban greening, are win-win-win solutions in all three positive nature futures (Fig. 4) to promote coexistence between humans and nature while minimising negative impacts on species and ecosystems (Karteris et al., 2016). As envisioned in our NFF narratives, Europe is moving towards renewable energy sources (Bórawski et al., 2019), in order to adapt to the European Climate Law (EC, 2023b). The expansion of renewable energy sources for Europe is essential to reduce net greenhouse gas emissions by at least 55% and reach carbon neutrality by 2050 (European Parliament, 2021). Urban greening is fundamental for human mental and physical health (Lee & Maheswaran, 2011) and for recreational and aesthetic appreciation (Veerkamp et al., 2021). Enhancing green areas is also relevant for cooling down cities, mitigating the effects of climate change, and reducing air pollution (Pauleit et al., 2020; Veerkamp et al., 2021). Community-based renewable energy and sustainable urban planning including zero-emission transportation, are examples of how to contribute to environmental sustainability, ecological connectivity, and improved human health simultaneously (Kammen & Sunter, 2016).

Our NFF narratives are adapted to the European context, but consistent with the interpretation given to the same framework in other studies (Pearson 2016; O’Connor et al., 2021). However, compared with other narratives developed at global scale (Pereira et al., 2020), in Europe the NaC perspective did not just focus on the relational value assigned to certain areas —such as the UNESCO Man and Biosphere reserves (MAB) (Reed, 2019)—, but also considered the historical value behind traditional practices associated with the European landscapes, such as vineyards or olive groves (UNESCO, 2014) and European Heritage sites (EC, 2024).

Narratives can be transformed into scenarios for environmental assessments, which are recognised as powerful tools for exploring how different pathways of societal development and policy choices could impact nature and the provision of NCP (Pereira et al., 2020). Some land-use and biodiversity models have been explored to determine whether it is possible to bend the biodiversity loss curve (Mace et al., 2018, Leclère et al., 2020). Although some scenarios demonstrated the feasibility of a positive outcome in this sense, there are still some limitations due to the challenges of further loss in several biodiversity-rich regions and threats, such as climate change, that have not been addressed (Pereira et al. in press). NFF scenarios provide more flexibility than previous ones, as they can reflect diverse values and worldviews, which helps identify context-relevant interventions (Kim et al., 2022). This has been done in Europe through scenario simulations which analyse synergies and trade-offs in land systems based on different value perspectives (Dou et al., 2023).

Our narratives can be interpreted and used as an additional layer that provides nuance and a representation of diversity in human-nature relational values to complement the macroeconomic assumptions of the SSPs/RCPs framework. At the same time, the development of these scenarios is a step towards revising the commonly used set of SSPs dominantly based on assumptions related to climate change mitigation and adaptation efforts, with nature playing a central role alongside existing socioeconomic considerations (Rosa et al., 2017).

Narratives can serve as the foundation for exploring the integration of land use and nature conservation scenarios to achieve the global biodiversity strategy goals (Pereira et al., 2020; Kim et al., 2023), and in the perspective of policy design in Europe, to achieve EU conservation goals for 2030. Systematic conservation planning (SCP) has been used to identify areas of conservation and restoration priorities for people and nature at both global (Strassburg et al., 2020; Jung et al., 2021) and EU (O’Connor et al., 2021) levels. Our NFF narratives can therefore be translated in settings for land use modelling and SCP and used as inputs for identifying opportunities and constraints for conservation and restoration in Europe. It may inform ongoing and upcoming conservation planning research, such as the achievement of the TEN-N (EC, 2020a), complementing the existing EU PA network in terms of species, habitats, and NCP, and to select suitable habitats within the future distributions of species and ecosystems in Europe.

Concerning the engagement process, the involvement of scientists with expertise in different fields offers the advantage of addressing all the topics covered by the narratives and spurring the ability of research to take different perspectives into account. However, the approach we adopted, especially accommodated visions and points of view of the conservation sector. For this reason, this imbalance may have skewed the interpretation of nature’s futures, lacking perspectives from diverse fields. The lack of participation of industry stakeholders may reflect low interest in the matter, a possible result of unawareness of their importance for achieving conservation objectives (Sterling et al., 2017). Indeed, this is something expected, because people that had a lower level of interest, as we highlighted in the mapping exercise, did not participate in the workshop. To address this challenge and solve issues concerning the process, some specific measures can be taken into account. To make the participatory process more balanced, efforts were made to address gaps emerged during the workshop by organising a post-workshop webinar to include a broader cross-section of society from different fields.

Overall, the communication between diverse fields may be convoluted due to sector-specific terminology, leading to varying interpretations of the discussions. To enhance the communication among stakeholders with different backgrounds, workshop notes were shared with all participants after the in-person workshop. Prior to the webinar, information on aims, NFF key-concepts, and technical terminology was provided to registered participants to facilitate their participation and contribution to the webinar.

While our narratives reveal the need for a more inclusive participatory process, the co-design approach carried out by conservationists envisages more constructive and preventive measures for nature, reflecting a more positive coexistence between humans and nature, which can be useful to model future scenarios and better steer EU policies towards the achievement of the 2030 conservation goals.

## Acknowledgements

We would like to acknowledge funding under the Horizon Europe project “NaturaConnect” (2022-2026). NaturaConnect receives funding under the European Union’s Horizon Europe research and innovation programme under grant agreement number 101060429. CR acknowledges additional funding by the European Union -NextGenerationEU. We would like to thank the NaturaConnect colleagues, but also the 13 stakeholders involved in the workshop and the 115 participants of the online event for their valuable participation and their contribution, which was fundamental in realising this study. Special thanks to Gabriele Rada who elaborated the illustrations of the nature futures narratives.

## Authors contribution

Carlo Rondinini, Peter H. Verburg, Henrique M. Pereira, Piero Visconti, Nestor Fernandez, Claudia Fornarini and Alessandra D’Alessio conceived the ideas and designed methodology; they also contributed in collecting the information during the stakeholders’ events together with Anandi Sarita Namasivayam, Jeremy Dertien, Martin Jung, Francisco Moreira, Louise O’Connor, Laura C. Quintero-Uribe, Martina Marei Viti; all authors contributed to write, edit and review the drafts. Alessandra D’Alessio and Claudia Fornarini led the writing of the manuscript. All authors gave final approval for publication.

## Conflict of Interest

The authors declare they do not have any conflicts of interest regarding the article.

## Notes

### Competing Interest Statement

The authors have declared no competing interest.

## References

Alexander, P., Henry, R., Rabin, S., Arneth, A., & Rounsevell, M. (2023). Mapping the shared socio-economic pathways onto the Nature Futures Framework at the global scale. Sustainability Science, 1–18. 10.1007/s11625-023-01415-z

Balmford, A. (2021). Concentrating vs. spreading our footprint: how to meet humanity’s needs at least cost to nature. Journal of Zoology, 315(2), 79–109. 10.1111/jzo.12920

Berbeć, A. K., Beata Feledyn-Szewczyk, B. (2018). Biodiversity of weeds and soil seed bank in organic and conventional farming systems. Institute of Soil Science and Plant Cultivation, State Research Institute in Puławy, Poland. 10.22616/rrd.24.2018.045

Bórawski, P., Bełdycka-Bórawska, A., Szymańska, E. J., Jankowski, K. J., Dubis, B., & Dunn, J. W. (2019). Development of renewable energy sources market and biofuels in The European Union. Journal of cleaner production, 228, 467–484. 10.1016/j.jclepro.2019.04.242

Brown, J. (2010). The World Café: Shaping our future through conversations that matter. www.ReadHowYouWant.com accessed on 27th October 2023.

Bugalho, M. N., Lecomte, X., Gonçalves, M., Caldeira, M. C., & Branco, M. (2011). Establishing grazing and grazing-excluded patches increases plant and invertebrate diversity in a Mediterranean oak woodland. Forest Ecology and Management, 261(11), 2133–2139. 10.1016/j.foreco.2011.03.009

Cannon, P. G., Gilroy, J. J., Tobias, J. A., Anderson, A., Haugaasen, T., & Edwards, D. P. (2019). Land-sparing agriculture sustains higher levels of avian functional diversity than land sharing. Global Change Biology, 25(5), 1576–1590. 10.1111/gcb.14601

Carvalho Ribeiro, S., Migliozzi, A., Incerti, G., & Pinto Correia, T. (2013). Placing land cover pattern preferences on the map: Bridging methodological approaches of landscape preference surveys and spatial pattern analysis. Landscape and Urban Planning, 114, 53–68. 10.1016/j.landurbplan.2013.02.011

CBD. POST-2020 GLOBAL BIODIVERSITY FRAMEWORK Draft recommendation submitted by the Co-Chairs. UN Doc Symbol CBD/WG2020/5/L.2 (5 December 2022).

Chan, K. M., Balvanera, P., Benessaiah, K., Chapman, M., Díaz, S., Gómez-Baggethun, E., … & Turner, N. (2016). Why protect nature? Rethinking values and the environment. Proceedings of the national academy of sciences, 113(6), 1462–1465. 10.1073/pnas.1525002113

Daskalova, G. N., & Kamp, J. (2023). Abandoning land transforms biodiversity. Science, 380(6645), 581–583. 10.1126/science.adf1099

Despotović, A., Joksimović, M., & Jovanović, M. (2020). Demographic revitalization of montenegrin rural areas through the smart village concept. Poljoprivreda i Sumarstvo, 66(4), 125–138. 10.17707/AgricultForest.66.4.10

Diez, J. J. & García, J. M., (2012).Sustainable forest management: An introduction and overview. Sustainable Forest Management: Current Research

Dotson, T., & Pereira, H. M. (2022). From antagonistic conservation to biodiversity democracy in rewilding. One Earth, 5(5), 466–469. 10.1016/j.oneear.2022.04.014

Dou, Y., Zagaria, C., O’Connor, L., Thuiller, W., & Verburg, P. H. (2023). Using the Nature Futures Framework as a lens for developing plural land use scenarios for Europe for 2050. Global Environmental Change, 83, 102766. 10.1016/j.gloenvcha.2023.102766

Dunn-Capper, R., Giergiczny, M., Fernández, N., Marder, F., & Pereira, H. M. (2024). Public preference for the rewilding framework: A choice experiment in the Oder Delta. People and Nature. 10.1002/pan3.10582

EC (2019). Report from the Commission to the European parliament, the council, the European economic and social committee and the committee of the regions. Review of progress on implementation of the EU green infrastructure strategy. COM 236. Brussels. https://eur-lex.europa.eu/legal-content/EN/TXT/PDF/?uri=CELEX:52019DC0236&qid=1562053537296&from=EN

EC (2016) No net land take by 2050? European Commission, Directorate-General for Environment, Publications Office of the European Union, 2016. https://data.europa.eu/doi/10.2779/537195.

EC (2020a). EU biodiversity strategy for 2030: bringing nature back into our lives. Communication from the commission to the european parliament, the council, the european economic and social committee and the committee of the regions. European Commission. COM 380. Brussels. https://data.europa.eu/doi/10.2779/048

EC (2020b). Report from the Commission to the European Parliament, the Council, the European Economic and Social Committee and the Committee of the Regions. The Farm to Fork Strategy for a fair, healthy and environmentally-Friendly food system. European Commission. COM 381. Brussels.

EC (2022a). Proposal for a Regulation of the European Parliament and of the Council on Nature Restoration. Brussels, 22.6. 2022, COM 304 Final 2022/0195 (COD).

EC (2023a). Energy communities. https://energy.ec.europa.eu/topics/markets-and-consumers/energy-communities_en accessed on 20th November 2023.

EC (2023b). European Climate Law. European Commission. https://climate.ec.europa.eu/eu-action/europeangreen-deal/european-climate-law_en accessed on 10th November 2023.

Edwards, D. P., Gilroy, J. J., Woodcock, P., Edwards, F. A., Larsen, T. H., Andrews, D. J., … & Wilcove, D. S. (2014). Land-sharing versus land-sparing logging: reconciling timber extraction with biodiversity conservation. Global change biology, 20(1), 183–191. 10.1111/gcb.12353

Grass, I., Batáry, P., & Tscharntke, T. (2021). Combining land-sparing and land-sharing in European landscapes. In Advances in Ecological Research (Vol. 64, pp. 251–303). Academic Press. 10.1016/bs.aecr.2020.09.002

Haga, C., Maeda, M., Hotta, W., Matsui, T., Nakaoka, M., Morimoto, J., … & Peterson, G. (2023). Modeling desirable futures at local scale by combining the nature futures framework and multi-objective optimization. Sustainability Science, 1–21. 10.1007/s11625-023-01301-8

Halada, L., Evans, D., Romao, C., & Petersen, J.-E. (2011). Which habitats of European importance depend on agricultural practices? Biodiversity and Conservation, 20(11), 2365–2378. https://10.1007/s10531-011-9989-z

IPBES (2016). The methodological assessment report on scenarios and models of biodiversity and ecosystem services. Ferrier, S., Ninan, K. N., Leadley, P., Alkemade, R., Acosta, L. A., Akçakaya, H. R., L., Brotons, W. W., Cheung, L., Christensen, V., Harhash, K. A., Kabubo-Mariara, J., Lundquist, C., Obersteiner, M., Pereira, H. M., Peterson, G., Pichs-Madruga, R., Ravindranath, N., Rondinini, C. and Wintle, B. A. (eds.)]. Secretariat of the Intergovernmental Science-Policy Platform on Biodiversity and Ecosystem Services, Bonn, Germany. 348 pages. 10.5281/zenodo.3235428

IPBES (2019). Global assessment report on biodiversity and ecosystem services of the Intergovernmental Science-Policy Platform on Biodiversity and Ecosystem Services.

Brondizio E. S., Settele J., Díaz S., Ngo H. T. (eds). IPBES secretariat, Bonn, Germany. 1148 pages. 10.5281/zenodo.3831673

IPBES (2023a). Scenarios and models. https://www.ipbes.net/scenarios-models accessed on 16th November 2023.

IPBES (2023b). Narrative Approaches. https://www.ipbes.net/narrative-approaches accessed on 16th November 2023.

Jung, M., Arnell, A., De Lamo, X., García-Rangel, S., Lewis, M., Mark, J., … & Visconti, P. (2021). Areas of global importance for conserving terrestrial biodiversity, carbon and water. Nature Ecology & Evolution, 5(11), 1499–1509. 10.1038/s41559-021-01528-7

Kammen, D. M., & Sunter, D. A. (2016). City-integrated renewable energy for urban sustainability. Science, 352(6288), 922–928. 10.1126/science.aad9302

Karteris, M., Theodoridou, I., Mallinis, G., Tsiros, E., & Karteris, A. (2016). Towards a green sustainable strategy for Mediterranean cities: Assessing the benefits of large-scale green roofs implementation in Thessaloniki, Northern Greece, using environmental modelling, GIS and very high spatial resolution remote sensing data. Renewable and Sustainable Energy Reviews, 58, 510–525. 10.1016/j.rser.2015.11.098

Kindlmann, P., & Burel, F. (2008). Connectivity measures: a review. Landscape ecology, 23, 879–890. 10.1007/s10980-008-9245-4

Kim, H., Peterson, G. D., Cheung, W. W. L., Ferrier, S., Alkemade, R., Arneth, A., Kuiper, J. J., Okayasu, S., Pereira, L., Acosta, L. A., Chaplin-Kramer, R., Den Belder, E., Eddy, T. D., Johnson, J. A., Karlsson-Vinkhuyzen, S., Kok, M. T. J., Leadley, P., Leclère, D., Lundquist, C. J., … Pereira, H. M. (2023). Towards a better future for biodiversity and people: Modelling Nature Futures. Global Environmental Change, 82, 102681. 10.1016/j.gloenvcha.2023.102681

Korpela, K., Borodulin, K., Neuvonen, M., Paronen, O., & Tyrväinen, L. (2014). Analyzing the mediators between nature-based outdoor recreation and emotional well-being. Journal of environmental psychology, 37, 1–7. 10.1016/j.jenvp.2013.11.003

Kremen, C. (2015). Reframing the land-sparing/land-sharing debate for biodiversity conservation. Annals of the New York Academy of Sciences, 1355(1), 52–76. 10.1111/nyas.12845

Kunming-Montreal Global biodiversity framework, 18 Dec. 2022, CBD/COP/15/L.25 [PDF-374 Kb]

Kuiper, J. J., Van Wijk, D., Mooij, W. M., Remme, R. P., Peterson, G. D., Karlsson-Vinkhuyzen, S., … & Pereira, L. M. (2022). Exploring desirable nature futures for Nationaal Park Hollandse Duinen. Ecosystems and People, 18(1), 329–347. 10.1080/26395916.2022.2065360

Kumar, R. S., Kundu, S., Kundu, B., Binu, N. K., & Shaji, M. (2021). Emerging typology and framing of climate-resilient agriculture in South Asia. In The Impacts of Climate Change (pp. 255–287). Elsevier. 10.1016/B978-0-12-822373-4.00021-5

Leclère, D., Obersteiner, M., Barrett, M., Butchart, S. H., Chaudhary, A., De Palma, A., … & Young, L. (2020). Bending the curve of terrestrial biodiversity needs an integrated strategy. Nature, 585(7826), 551–556. 10.1038/s41586-020-2705-y

Lee, A. C., & Maheswaran, R. (2011). The health benefits of urban green spaces: a review of the evidence. Journal of public health, 33(2), 212–222. 10.1093/pubmed/fdq068

Lindroth, A., Vestin, P., Sundqvist, E., Mölder, M., Bâth, A., Hellström, M., … & Weslien, P. (2012, April). Clear-cutting is causing large emissions of greenhouse gases-are there other harvest options that can avoid these emissions? In EGU general assembly conference abstracts (p. 7578).

Lundquist, C., Hashimoto, S., Denboba, M. A., Peterson, G., Pereira, L., & Armenteras, D. (2021). Operationalizing the Nature Futures Framework to catalyse the development of nature-future scenarios. Sustainability Science 16(6), 1773–1775. 10.1007/s11625-021-01014-w

Mace, G. M., Barrett, M., Burgess, N. D., Cornell, S. E., Freeman, R., Grooten, M., & Purvis, A. (2018). Aiming higher to bend the curve of biodiversity loss. Nature Sustainability, 1(9), 448–451. 10.1038/s41893-018-0130-0

Mansur, A. V., McDonald, R. I., Güneralp, B., Kim, H., de Oliveira, J. A. P., Callaghan, C. T., … & Pereira, H. M. (2022). Nature futures for the urban century: Integrating multiple values into urban management, Environmental Science & Policy, Vol. 131, Pag. 46-56. 10.1016/j.envsci.2022.01.013

O’Connor, L. M., Pollock, L. J., Renaud, J., Verhagen, W., Verburg, P. H., Lavorel, S., … & Thuiller, W. (2021). Balancing conservation priorities for nature and for people in Europe. Science, 372(6544), 856–860. 10.1126/science.abc4896

O’Neill B. C., Kriegler E., Riahi K., Ebi K., Hallegatte S., Carter T. R., Mathur R., van Vuuren, D. P. (2014). A new scenario framework for climate change research: The concept of shared socio-economic pathways. Climatic Change 122, 387–400. 10.1007/s10584-013-0905-2

Obermeister, N. (2019). Local knowledge, global ambitions: IPBES and the advent of multi-scale models and scenarios. Sustain Sci 14, 843–856. 10.1007/s11625-018-0616-8

Pascual, U., Balvanera, P., Anderson, C.B. et al. (2023). Diverse values of nature for sustainability. Nature 620, 813–823 . 10.1038/s41586-023-06406-9

Pauleit, S., Hansen, R., Rall, E. L., & Rolf, W. (2020). Urban green infrastructure: Strategic planning of urban green and blue for multiple benefits. In The Routledge Handbook of Urban Ecology (pp. 931–942). Routledge. 10.1016/j.jclepro.2019.04.242

Pearson, R. G. (2016). Reasons to conserve nature. Trends in Ecology & Evolution, 31(5), 366–371. 10.1016/j.tree.2016.02.005

Pereira, L. M., Davies, K. K., den Belder, E. Ferrier, S., Karlsson-Vinkhuyzen, S., Kim, H., …& Lundquist, C. J. (2020). Developing multiscale and integrative nature–people scenarios using the Nature Futures Framework. People Nat. 2: 1172– 1195. 10.1002/pan3.10146

Pereira, H. M., Rosa, I. M., Martins, I. S., Kim, H., Leadley, P., Popp, A., … & Alkemade, R. Global trends in biodiversity and ecosystem services from 1900 to 2050. . Science, in press.

Quintero-Uribe, L. C., Navarro, L. M., Pereira, H. M., & Fernández, N. (2022). Participatory scenarios for restoring European landscapes show a plurality of nature values. Ecography, 2022(4), e06292. 10.1111/ecog.06292

Renting, H., Rossing, W. A., Groot, J. C., Van der Ploeg, J. D., Laurent, C., Perraud, D., … & Van Ittersum, M. K. (2009). Exploring multifunctional agriculture. A review of conceptual approaches and prospects for an integrative transitional framework. Journal of environmental management, 90, S112–S123. 10.1016/j.jenvman.2008.11.014

Reed, M.G. The contributions of UNESCO Man and Biosphere Programme and biosphere reserves to the practice of sustainability science. Sustain Sci 14, 809–821 (2019). 10.1007/s11625-018-0603-0

NaturaConnect (2024). Deliverables, D5.1 Scenario framework for TEN-N, translation of NFF storylines into indicators and scenario settings. https://naturaconnect.eu/deliverables/

Rosa, I. M. D., Pereira, H. M., Ferrier, S., Alkemade, R., Acosta, L. A., Akcakaya, H. R., … & Van Vuuren, D. (2017). Multiscale scenarios for Nature futures. Nat Ecol Evol 1, 1416–1419. 10.1038/s41559-017-0273-9

Saito, O., Kamiyama, C., Hashimoto, S., Matsui, T., Shoyama, K., Kabaya, K., Uetake, T., Taki, H., Ishikawa, Y., Matsushita, K., Yamane, F., Hori, J., Ariga, T. & Takeuchi, K. (2019). Co-design of national scale future scenarios in Japan to predict and assess natural capital and ecosystem services. Sustain Sci 14, 5–21. 10.1007/s11625-018-0587-9

Scherer, L. A., Verburg, P. H., & Schulp, C. J. (2018). Opportunities for sustainable intensification in European agriculture. Global Environmental Change, 48, 43–55. Advance online publication. 10.1016/j.gloenvcha.2017.11.009

Schindler, S., Sebesvari, Z., Damm, C., Euller, K., Mauerhofer, V., Schneidergruber, A., … & Wrbka, T. (2014). Multifunctionality of floodplain landscapes: relating management options to ecosystem services. Landscape Ecology, 29, 229–244. 10.1007/s10980-014-9989-y

Secretariat of the Convention on Biological Diversity (2020). Global Biodiversity Outlook 5. Montreal

Sidemo-Holm, W., Ekroos, J., & Smith, H. G. (2021). Land sharing versus land sparing—What outcomes are compared between which land uses? Conservation Science and Practice, 3(11), e530. 10.1111/csp2.530

Sterling, E. J., Betley, E., Sigouin, A., Gomez, A., Toomey, A., Cullman, G., … & Porzecanski, A. L. (2017). Assessing the evidence for stakeholder engagement in biodiversity conservation. Biological conservation, 209, 159–171. 10.1016/j.biocon.2017.02.008

Strassburg, B. B., Iribarrem, A., Beyer, H. L., Cordeiro, C. L., Crouzeilles, R., Jakovac, C. C., … & Visconti, P. (2020). Global priority areas for ecosystem restoration. Nature, 586(7831), 724–729. 10.1038/s41586-020-2784-9

Tittensor, D. P., Walpole, M., Hill, S. L. L., Boyce, D. G., … & Ye, Y. (2014). A mid-term analysis of progress toward international biodiversity targets. Science 346, 241–244. 10.1126/science.1257484

Toader, M., Roman, G. V., & Năstase, P. I. (2018). Socio-economic and cultural revitalization of rural localities-an essential challenge of the Central and Eastern European countries. Lucrări Ştiinţifice – vol. 61(2), seria Agronomie. https://repository.uaiasi.ro/xmlui/handle/20.500.12811/545

UNESCO, SCBD. “Florence declaration on the links between biological and cultural diversity.” (2014).

van der Wal, R., Miller, D., Irvine, J., Fiorini, S., Amar, A., Yearley, S., … Dandy, N. (2014). The influence of information provision on people’s landscape preferences: A case study on understorey vegetation of deer-browsed woodlands. Landscape and Urban Planning, 124, 129–139. 10.1016/j.landurbplan.2014.01.009

Veerkamp, C. J., Schipper, A. M., Hedlund, K., Lazarova, T., Nordin, A., & Hanson, H. I. (2021). A review of studies assessing ecosystem services provided by urban green and blue infrastructure. Ecosystem Services, 52, 101367. 10.1016/j.ecoser.2021.101367

Zerbe, S. (2022). Restoration of Multifunctional Cultural Landscapes: Merging Tradition and Innovation for a Sustainable Future (Vol. 30). Springer Nature. 10.1007/978-3-030-95572-4

