## Supplementary material 1, Figure S1, Supplementary material 2, Figure S2, Figure S3, Supplementary material 3 for "Narratives for positive nature futures in Europe"

### Supporting Information

#### Supplementary material 1: Nature Futures Framework (NFF) triangle

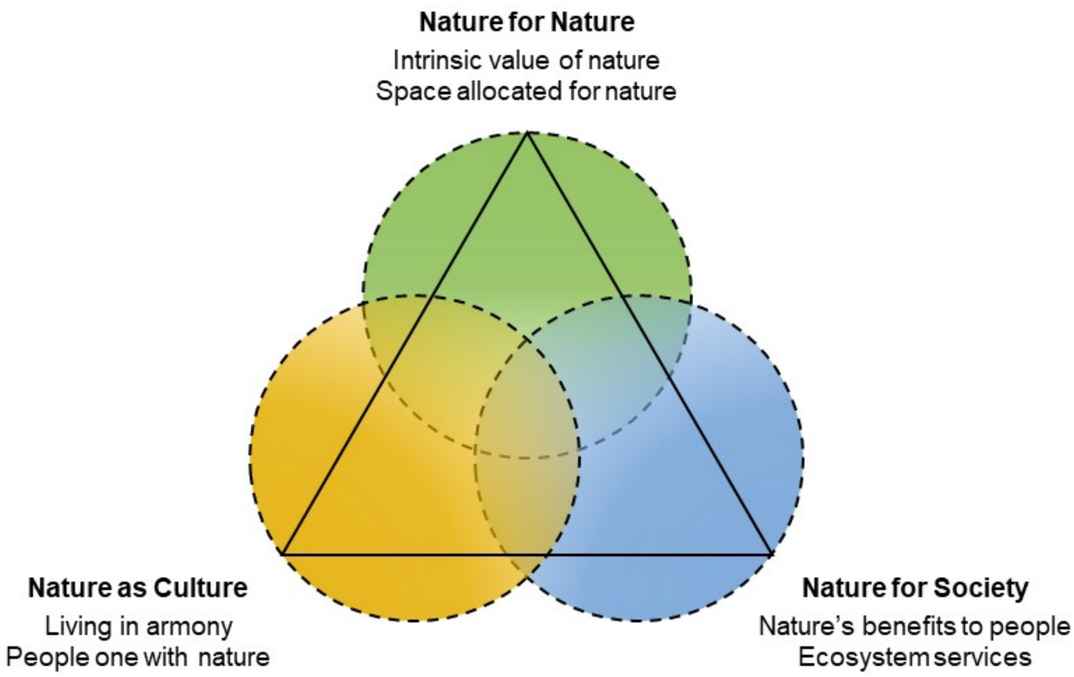


**Figure S1**: The Nature Futures Framework presents three value perspectives of nature in a triangle. Source: Adapted from Pereira et al., 2020.

#### Supplementary material 2: Workshop and webinar

##### Appendix 2. Supplementary materials and methods

The methodological approach consists of four main phases divided in ten steps that led to the elaboration of the draft narratives after the stakeholders' identification and the main engagement event. This was followed by the organisation of a second engagement event to fill the gaps, and the refinement of the narratives' final version through a second elicitation stage and the cross-narratives analysis.

The preparatory phase to the stakeholders elicitation consisted of *1)* identifying a set of EU macroeconomic, social and legislative assumptions, or 'constraints', that coerce the NFF narratives; *2*) formulating questions on nature futures based on the constraints and identifying a set of priority themes to be discussed with the stakeholders, and *3)* selecting stakeholder experts on different fields through a mapping exercise, which focused on their importance and influence across Europe.

The second phase was the first stakeholders' event i.e. *4)* an in-person workshop to elicit preferences around the themes and perspectives on the future of nature.

The third phase consisted of *5)* the elaboration of three draft narratives based on the outcomes of the workshop and *6)* the formulation of additional questions to be asked during a second stakeholders' event on nature futures.

During the fourth phase, *7)* the questions were presented during the second engagement stage which was an online event, to obtain feedback on the draft narratives and fill potential gaps; *8)* Eventually, the narratives were then refined integrating the outcomes into a second set of draft narratives, and *9)* further reviewed by domain experts, which allowed us *10)* to develop the final set of narratives following the Nature Futures Framework.

#### Appendix 2.1 Preparatory phase to the stakeholders' elicitation: identification of constraints and preliminary themes for the narrative, and formulation of broad questions:

In order to pinpoint the constraints that coerce the narratives (**step 1**) we examined the EU legislation, regulations, goals, and strategic priorities that are essential for NFF narratives.We took into account the following key strategic goals:

- the increase in the coverage of protected areas (PAs) to a minimum of 30% for both land and sea, including 10% under strict protection. This expansion is intended to establish sufficiently large areas where vital natural processes can occur undisturbed (EC, 2022a). It also foresees the enhancement of the Natura 2000 sites and nationally PAs, improving the conservation status of species and habitats, and contributing to addressing future environmental changes (EEA, 2020);
- the European Nature Restoration Law (NRL), which targets the restoration of 20% of EU land. This initiative supports the recovery of ecosystems and species towards optimal ecological conditions (EC, 2022a). It also includes specific restoration actions for pollinators, river connectivity, forests, agriculture, urban areas, and marine ecosystems. Objectives encompass reversing the decline of pollinator populations, restoring rivers to a free-flowing state, achieving no net loss of green urban space by 2030, and increasing biodiversity in agricultural landscapes (EC, 2022a);
- the global goal of "No Net Land Take" by 2050, regarding the urban areas system. Assuming a linear progression in land take, achieving the EU's goal of reaching zero net land take by 2050 needs an annual reduction of 14 km2 from 2019 onward.

Consequently, by 2030, the EU must decrease its yearly net land take to 282 km2; (EC, 2016)

- the EU Farm to Fork Strategy goals which were essential to set the constraints for the agriculture context. Such goals include transitioning 25% of agriculture to organic farming, reducing chemical pesticide use by 50%, and decreasing fertiliser use by 20% by 2030 (EC, 2020a);
- the European Climate Law (Regulation (EU) 2021/1119) that commits Member States to achieve the EU's climate goal of reducing emissions by at least 55% by 2030 compared to 1990. This involves measures such as increasing renewable energy sources to 32%, reducing fossil fuel biomass, planting three billion trees, and restoring carbon-rich ecosystems by 2030 (EC, 2023b).

Considering the main EU policy objectives, we examined a set of themes that are crucial for the future of nature and challenging for its conservation (**step 2**). After a brainstorming, which involved different domain experts, the following themes emerged: urban systems, forestry, freshwater ecosystems, green and blue infrastructure, habitats conservation and ecosystem restoration, agroecological policies, infrastructure development and renewable energies, and species conservation. We then formulated questions on the above mentioned themes, but also on specific priorities in PAs planning and connectivity design, to address stakeholders' visions of positive futures for people and nature in Europe, according to the different perspectives of the NFF. Some examples were: 'What are the main changes happening in the landscapes in this Nature Futures (NF)?', 'What are the dominant changes in the management of agricultural areas?', 'Why do people conserve nature according to this NF vision?', 'What type of Nature-based Solutions do you expect according to this NF vision?'

#### Appendix 2.2. Identification of key stakeholders

The initial step of any stakeholder engagement is the mapping of stakeholder groups according to their perceived influence and interest in the selected subject.

The mapping exercise was conducted with a power-engagement grid, which organises them into four groups based on their interest in the process and their influence on its outcomes (Fig. S2).

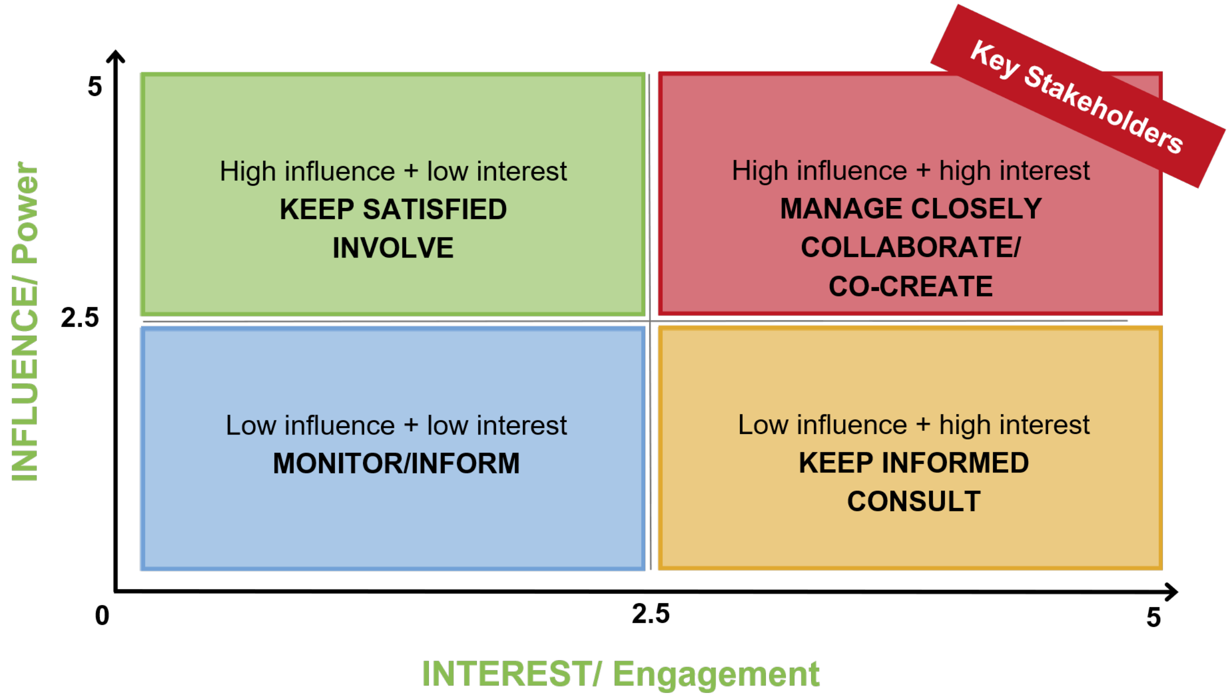


**Figure S2**: The conceptual framework of the Mendelow Matrix was applied by NaturaConnect experts leading the work on the NFF, to identify plausible and supported Nature Future narratives that are compatible with the achievement of the objectives of the EU Biodiversity Strategy 2030. The axes show INFLUENCE/Power and INTEREST/Engagement values in a scale from 0 to 5.  © NaturaConnect/EUROPARC Federation, adopted from Mendelow (1981).

This exercise helps determine the right level of engagement for each stakeholder. For example, key players with high power and high interest should be deeply involved in co-design processes, while stakeholders with high power but low interest should be kept engaged at an appropriate level to keep them satisfied and informed, ideally raising their interest over time to engage them at a later stage. Identified stakeholder groups include European policy and decision makers (e.g. EU commissions, EU Member State representatives and authorities), research institutes and national and international NGOs. To understand stakeholders perspectives on designing future biodiversity protection in Europe, relevant sectors and interest groups were identified. Although we contacted stakeholders from across Europe, we placed particular focus on representatives from the NaturaConnect case study areas (Finland, Portugal, France, Leipzig-Halle and Doñana). The identified stakeholders, whose list can be found in Tables S1 and S2, were then invited to participate in person to a workshop held in Leipzig (step 4)

##### Appendix 2.3 First elicitation stage: workshop to address questions and elicit stakeholders' visions

The broad questions were asked during the first stakeholder engagement event, which was a three-day workshop, held in Leipzig (Germany) from 8 to 10 May 2023 (**step 4**). At this event, the scientists affiliated with the NaturaConnect project and with specific expertise were introduced as internal stakeholders.

Considering the major importance of PAs planning and connectivity, we decided to dedicate Day 1 to the elicitation of diverse visions of NF for Europe, Day 2 to connectivity design, and Day 3 to planning PAs in the context of different NF.

We adopted the Appreciative Inquiry methodology as a method to facilitate high levels of stakeholder engagement. This approach encourages stakeholders to explore existing successes, identify needs, and support organisational and sectoral change. It emphasises reflective questioning and focuses on envisioning future possibilities rather than just solving current problems (Armstrong, Holmes & Henning 2020).  During the  morning session of Day 1, the NFF was presented to the participants, placing specific emphasis on highlighting the distinctions among the three nature futures perspectives. Following this, participants were asked to envision the future of European landscapes from the three perspectives. We asked participants to borrow some pictures of European landscapes which they were attached to and towards which they felt the need to reflect on. In addition, we showed different pictures of European landscapes related to the themes previously identified, to facilitate the discussions and elicit the stakeholders' preferences about them, such as the configuration of urban systems or the renewable energy expansion  (Fig. S3).

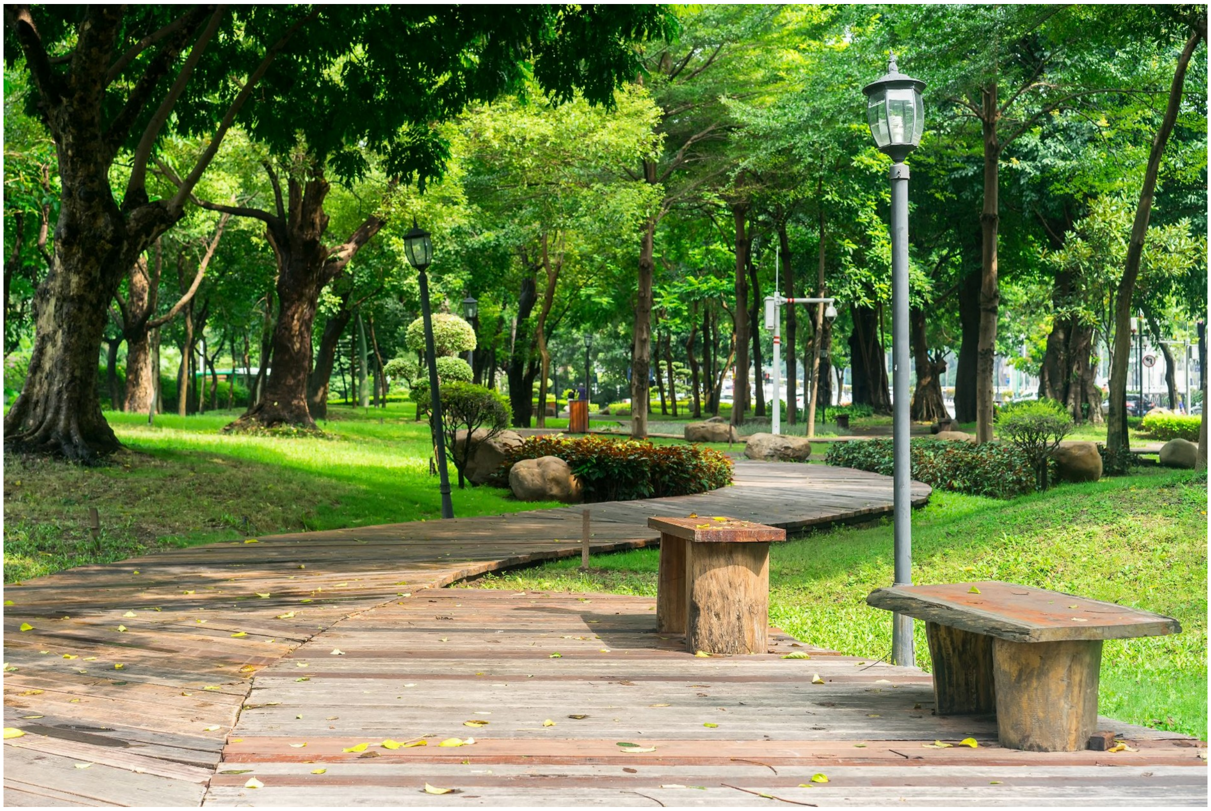

1

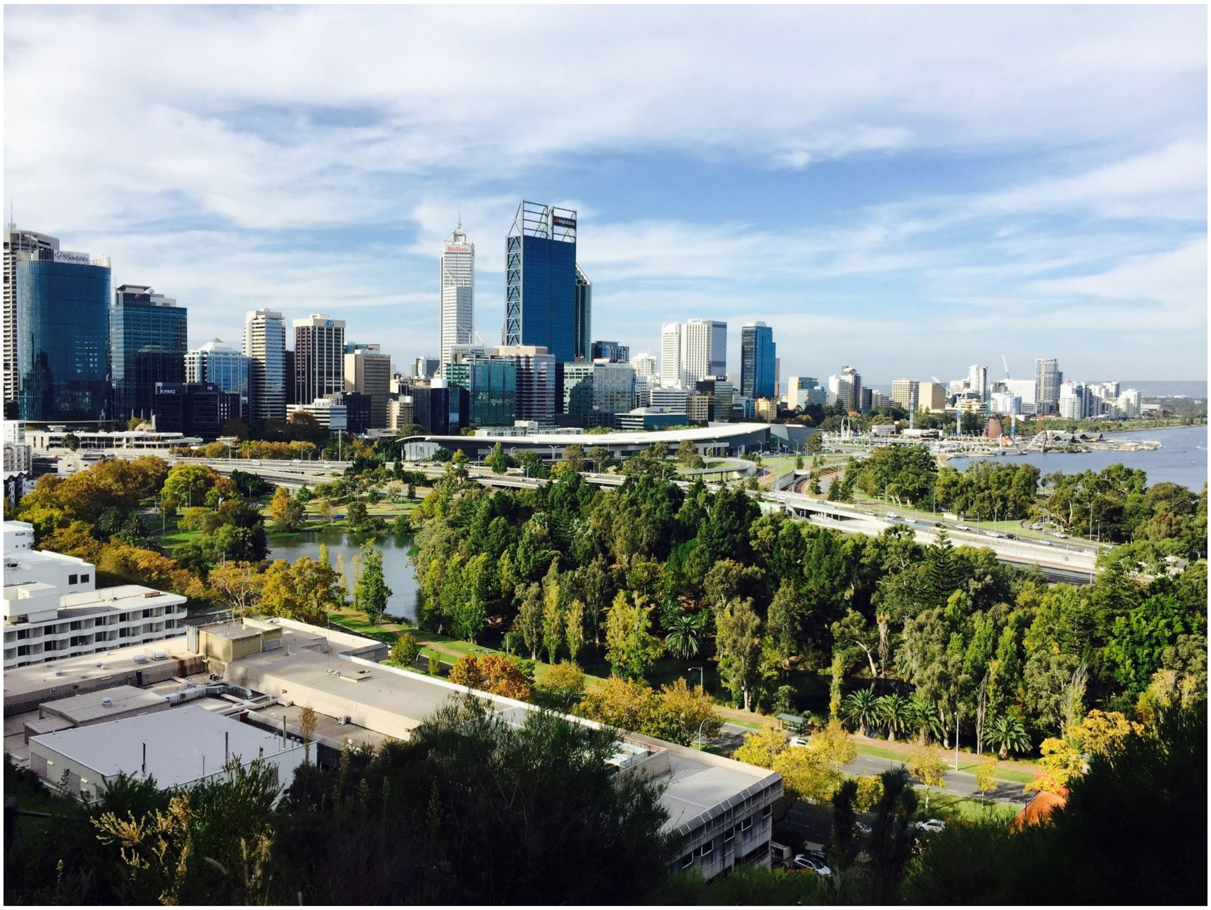

1

1

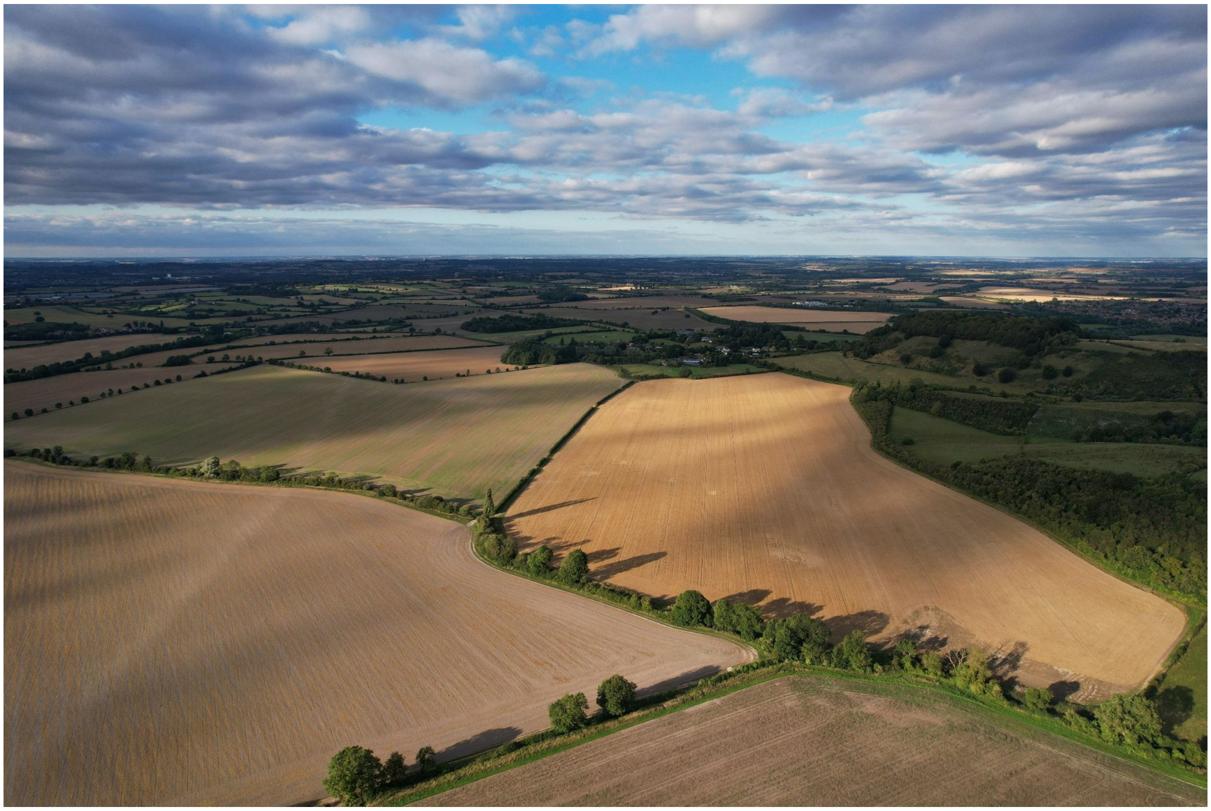

1

1

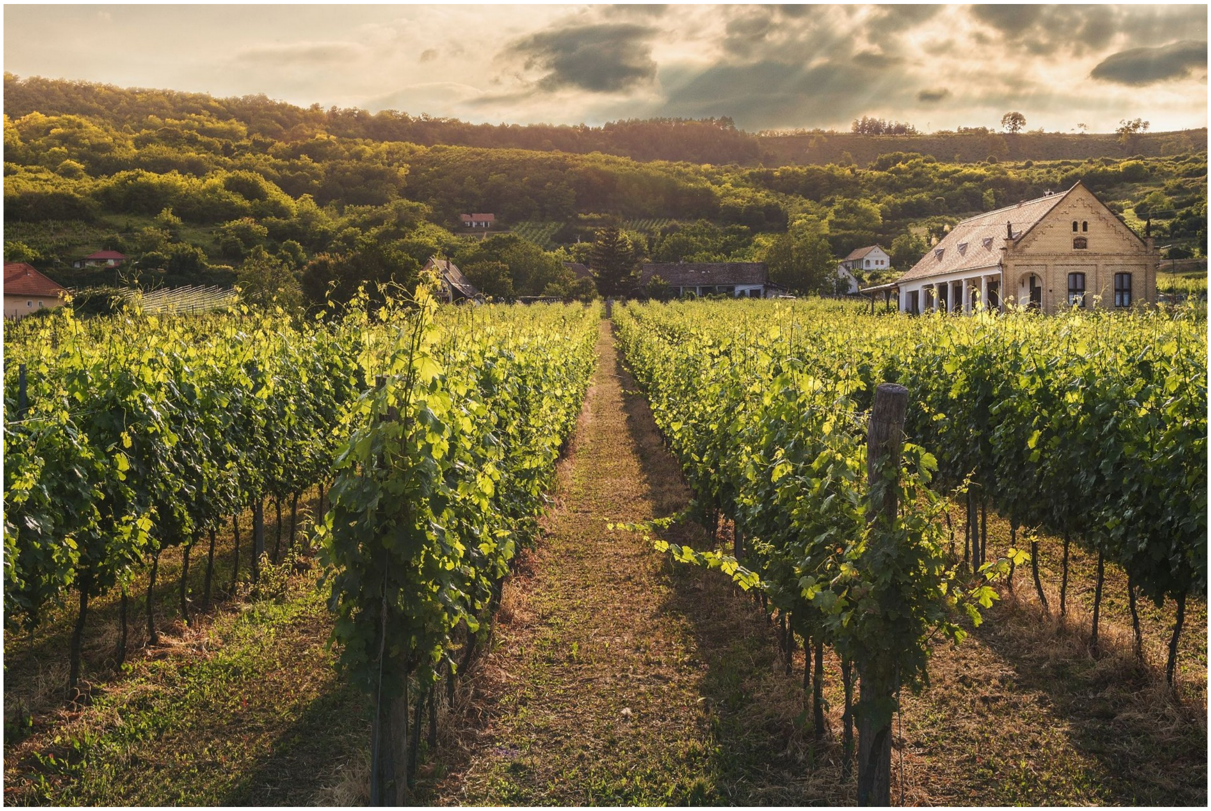

1

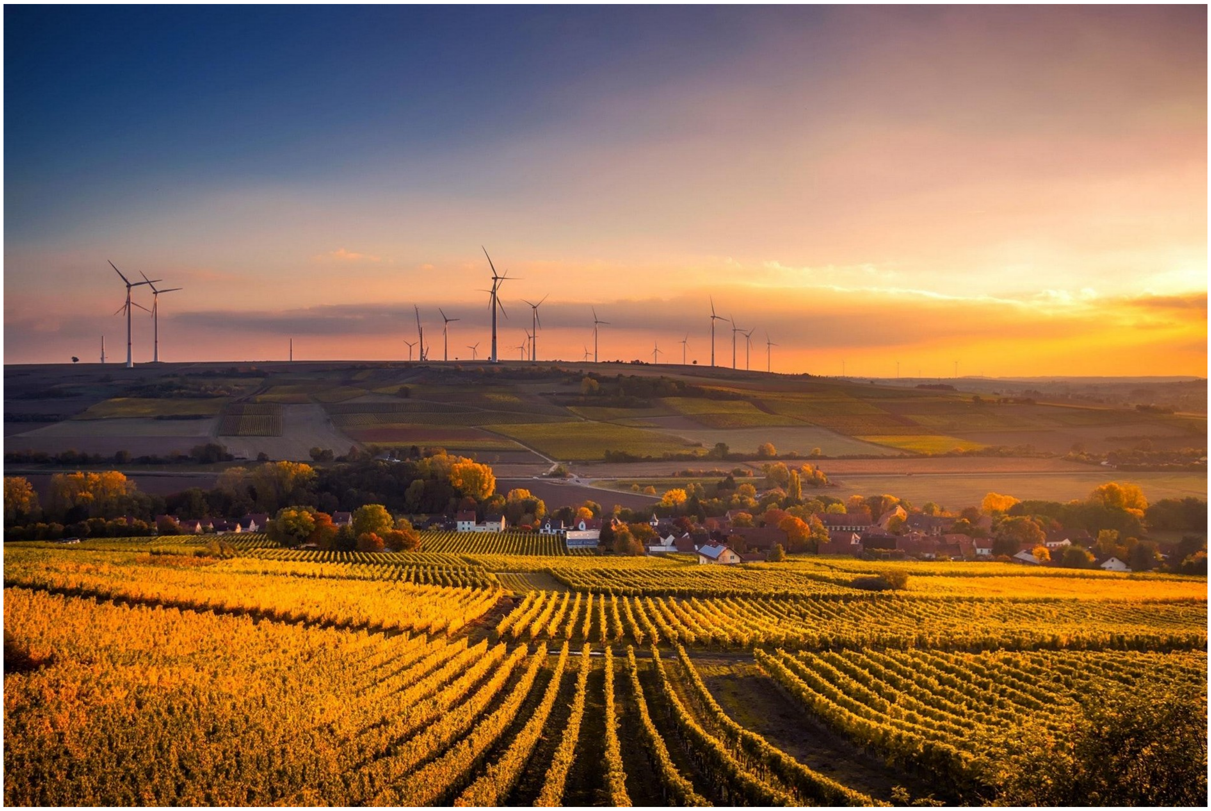

2

3

1

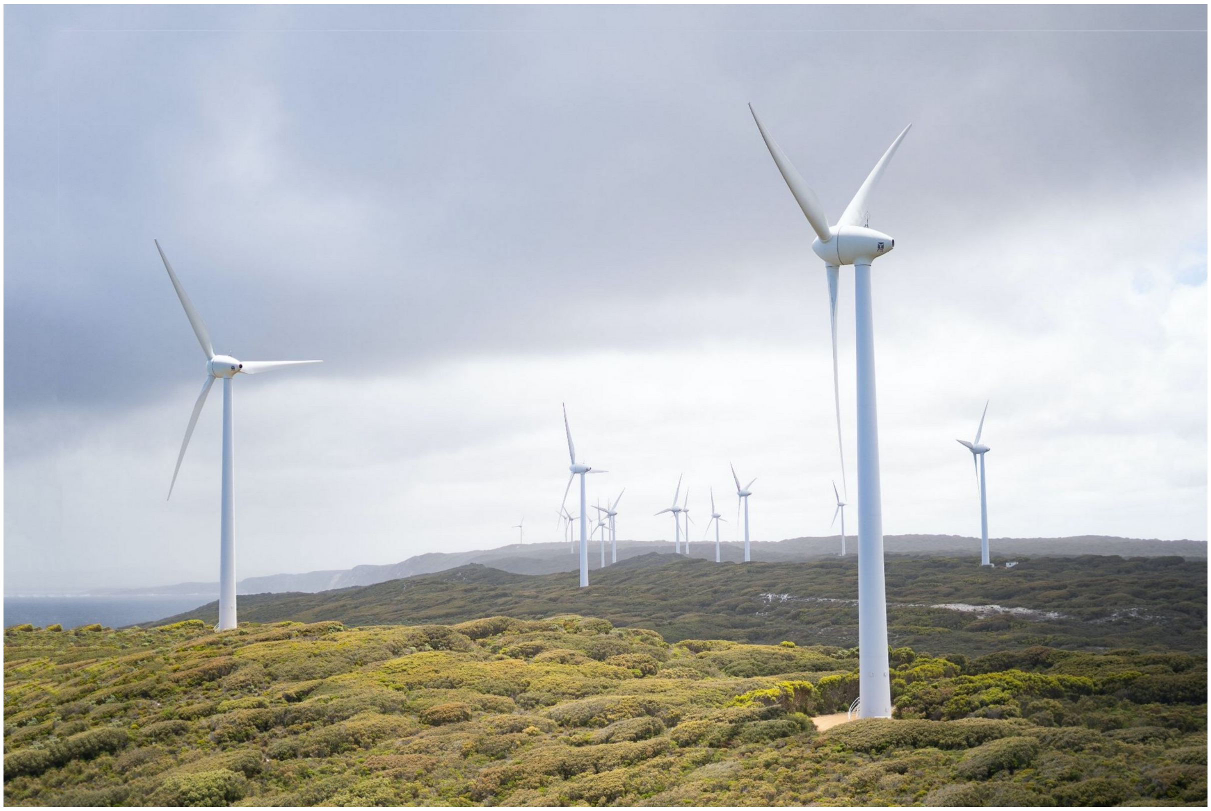

1

2

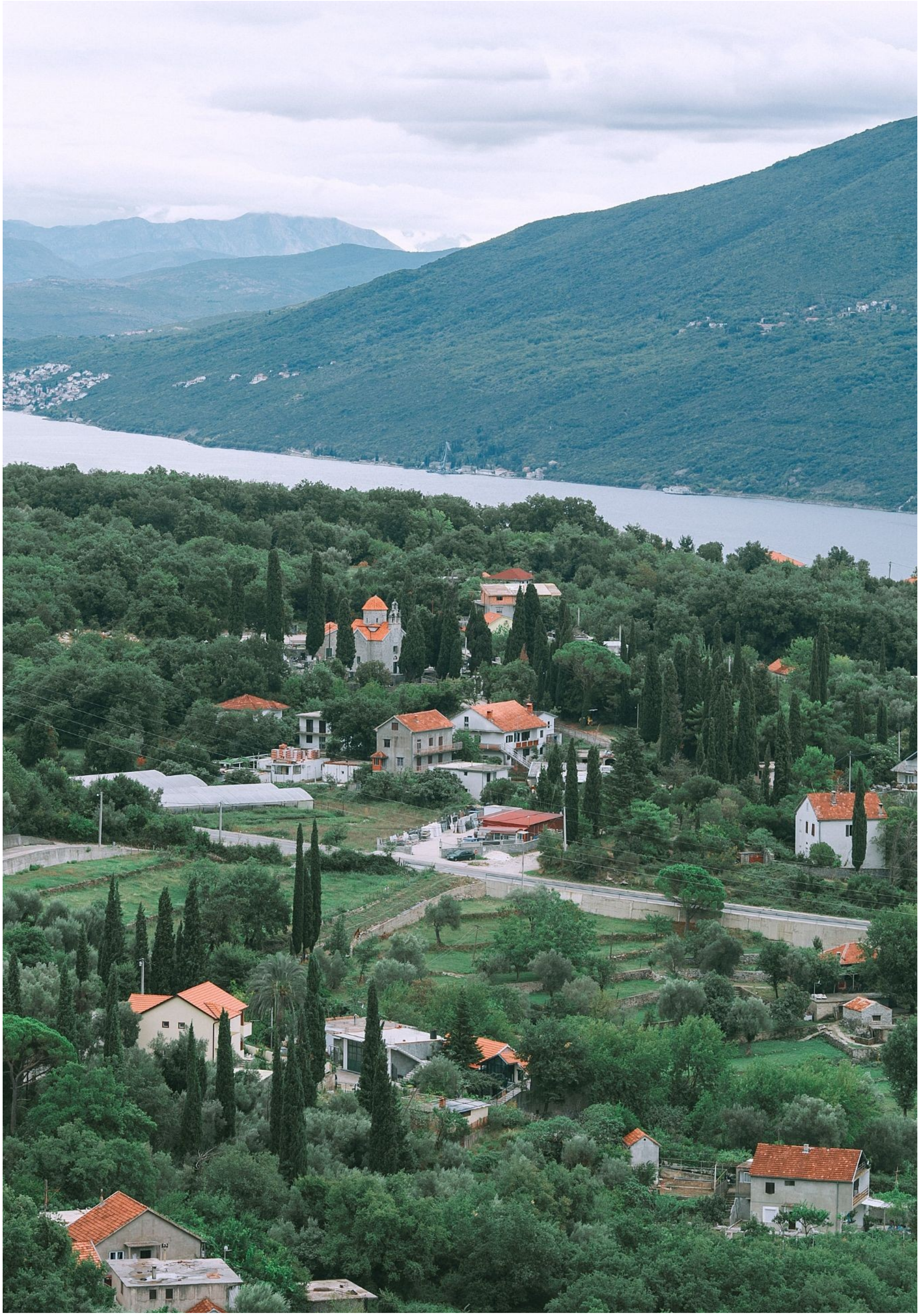

**Figure S3**: Pictures of different European landscapes shown on the first day of the workshop during the first round of the World Café Method. © Pexels

We specifically adopted the World Café engagement method, a structured technique to foster open dialogue and exchange of various ideas, allowing people to move among tables of conversation (Brown, 2010). It generally starts creating a welcoming and relaxed atmosphere, resembling a café, with small tables. Participants gather at tables to discuss a specific question or topic for about 20-30 minutes per round. After each round, they move to different tables, with one "table host" staying behind to summarise the previous conversation for the new group (Brown, 2010). In this case, we organised people into three groups, each corresponding to one of the NF corners and arranged multiple rounds of 30-minute conversation. Afterwards, each participant moved among tables, contributing to different discussions covering all aspects of the three corners of the NFF triangle. Since the NFF and related discussion may be something difficult to understand for people not familiar with it, we decided to keep an internal stakeholder as a moderator at each table to facilitate the conversation and guide the dialogues. The moderators also prompted the discussion with the questions and took notes of the answers. In addition, to make each person actively involved in the discussion we provided boards, post-it notes and visual items. In this way, even if one person did not take the floor during the discussion, we ensured that written ideas and reflection were collected and incorporated in the outcomes.

During the afternoon session on Day 1, attendees shared their perspectives on the themes previously identified (Appendix 2.1). Employing the World Café method again, the session facilitated multiple rounds of conversation, enabling participants to self-organise and engage in discussions aligned with their interests around five tables, one per each topic.

On Day 2, the focus was on connectivity. The initial discussion utilised the World Café format with six tables, featuring two tables per NF corner, and it was facilitated by moderators who took notes on participants' answers to specific questions. Each table addressed three primary questions: 'Why is connectivity important in this NF?', 'What are the main threats to preserving and enhancing connectivity?', and 'What species, ecological processes, and/or ecosystem services should be prioritised?' Participants rotated between tables every 20 minutes, engaging in diverse conversations across different corners. A subsequent set of questions included: 'What type of areas do we need to connect in this NF?', 'Where should we allocate connectivity?', and 'What will be the impact of developing other infrastructures?'.

Following this step, a collective brainstorming session was conducted to emphasise priorities, enablers, and obstacles in implementing connectivity. Participants documented their ideas on five distinct boards, each associated with one policy framework or activity sector: Green and Blue Infrastructure, habitat conservation and ecosystem restoration, agroecological policies, infrastructure development and renewable energies, and species conservation. Participants were given the autonomy to choose their own topic and time for discussion.

Day 3 delved into aspects of protected areas planning through two rounds of conversation. Participants discussed the same set of questions in five tables together with moderators, aiming to highlight differences among the three value perspectives. The four main questions included: 'Where should strict protection take place and why, in each NF?', 'What should take priority in each NF in terms of identifying and managing the rest of the Protected Area (PA) network?', 'Where and why would you allow some human activities inside PAs?', and 'Where and why would you have larger or smaller PAs in each NF corner?)'. During the afternoon of Day 2 and Day 3, discussions were not addressed towards the three NFF perspectives.

Immediately after the workshop, the notes taken by each moderator were collected, cleaned up and organised in a shared document, based on the days of the workshop and the topic they referred to. Thus, the folder with all the notes was shared to the participants, who were asked to add further input if considered worthwhile.

##### Appendix 2.3 Elaboration of draft narratives based on the workshop outcomes and formulation of further specific questions

Shortly after the workshop, a series of meetings and interviews were conducted with all moderators to ensure a comprehensive understanding of the discussions at each table and develop the first set of the narratives (**step 5**) . The goal was to further refine the notes gathered during the workshop. Subsequently, all the inputs provided by participants were thoroughly checked and organised according to the NFF corners. In addition, we streamlined the narratives, narrowing down the focus to a concise selection of recurring topics, including Urban systems, Forestry, Freshwater Ecosystems, and Energy. We also synthesised agro-ecological policies under the main topic of Agriculture, and grouped the other themes and the outcomes of Day 2 and Day 3 under Nature Protection and Restoration. Thus, we focused the narratives on this smaller set of six topics. By doing the analysis, gaps and inconsistencies among the narratives emerged,  particularly on the preferences related to Nature and Restoration, and Agriculture. To gather feedback and obtain additional insights on the current description of these narratives, a review process was initiated, together with the NaturaConnect consortium. We formulated further specific questions to be asked during a second stakeholders' engagement event (**step 6**):

1) In a Nature for Nature scenario, what activities would you restrict in strictly protected areas?
2) In a Nature for Society scenario, what activities would you restrict in strictly protected areas?
3) In a Nature as Culture scenario, what activities would you restrict in strictly protected areas?
4) In a Nature for Nature scenario, what kinds of forestry activities would be allowed?
5) In a Nature for Society scenario, what kinds of forestry activities would be allowed?
6) In a Nature as Culture scenario, what kinds of forestry activities would be allowed?
7) In a Nature for Nature scenario, what types of agricultural land uses should be promoted?
8) In a Nature for Society scenario, what types of ecosystem services can be reinforced in agricultural landscapes?
9) In a Nature as Culture scenario, what cultural landscapes are important for nature conservation?
10) How important is the reduction of agricultural land in each scenario?
11) In which scenario(s) do you think high-density urban areas should be emphasised?
12) Which green elements should be integrated into which scenarios?
13) In a Nature for Nature scenario, in which areas and ecosystems should we implement large scale rewilding?
14) In a Nature for Society scenario, where could ecological corridors be prioritised?
15) In a Nature as Culture scenario, what measures can contribute to improve Green Infrastructure?

#### Appendix 2.4 Refinement of the final narratives version: second elicitation stage

A 2-hour online public webinar titled 'Nature Future Scenarios for a Resilient Trans-European Nature Network (TEN-N)' was held on July 4, 2023 (**step 7**). The event was mainly advertised online by the NaturaConnect consortium, via websites and social media. However, to ensure a heterogeneous group of stakeholders and engage experts of specific sectors missing from the previous event, e.g. for example those that were interested but unable to commit to a multi-day in person workshop, we invited additional key stakeholders identified from the previous mapping exercise, and opened up the engagement to a greater representation. 115 people joined the webinar, however it was not conclusively possible to determine their field of expertise, since only 60% participants answered the questions about the sector they belong to. The draft narratives for each topic were introduced by scientists from the NaturaConnect consortium, highlighting the contrasting perspectives within the three narratives. Following each presentation, participants were engaged in Q&A sessions, responding via Mentimeter (https://www.mentimeter.com/). The Mentimeter questions, designed to gather additional input and address gaps that were identified in the in-person workshop, aimed to enhance the narratives. Post-webinar, the collected responses were analysed, identifying recurring statements and integrating participant feedback into the narratives (**step 8**). The entire webinar, including discussions and presentations, was recorded and made publicly accessible as an online resource. The second version of the narratives were reviewed by the NaturaConnect consortium member and moderators of the stakeholder events (**step 9**). This step ensured that the statements in the revisited narratives relied on the initial stakeholders' inputs and were realistic within the European context.

Finally, we integrated the reviews and produced the final version of the narratives (**step 10**).

### Supplementary material 3: Supplementary discussion

#### **Appendix 3.** Supplementary discussion on the differences among the narratives

Agriculture is one of the major drivers of biodiversity loss with increasing impacts due to changes in consumption patterns and growing populations (Chaudhary et al., 2016; Dudley & Alexander, 2017). Protection of biodiversity is indeed a prerogative in modern agriculture, which is reflected at the European level in the EU Biodiversity Strategy (EC, 2020a) that aims to reverse the loss of biodiversity and Nature's Contributions to People (NCP) in the member states by 2030 (Berbeć & Feledyn-Szewczyk, 2018). In NfS, agricultural production is expected to take place in highly productive areas not of conservation concern, also giving priority to regulating services. Therefore, the species and varieties richness of both cultivated and wild plants, livestock and wild animals of rural areas must be protected for maintaining ecological services that provide soil fertility and productivity of agricultural ecosystems (Clergue et al., 2005; Feledyn-Szewczyk & Berbeć, 2018). In a Nature for Nature (NfN) perspective large-scale farming is envisioned except for areas within and next to protected areas. This method is related to the land sparing approach (Fischer et al., 2008) to face the challenge of feeding the growing human populations (Marcacci et al., 2020), which has led to changes in agricultural practices over time with the development of new technologies, the enhancement of crop varieties and the use of agrochemicals (Pingali, 2012). In this context, Nature Based Solutions (NBS) such as integrated pest management, regenerative farming and precision farming are envisioned. Even if excessive costs and lack of incentive policies, may limit the spread of these latter technique (Troiano et al., 2023), in both NfN and Nature for Society (NfS) they all mitigate the effects of pesticide use and chemical inputs, provide minimum natural elements in the landscapes such as woodland islets and hedgerows, stone walls, etc. and optimise both agricultural input and output while reducing extra water consumption during irrigation (Finger et al., 2019; Raj et al., 2021). Contrary to the NfN scenario, in Nature as Culture (NaC) large-scale farming gets converted to small-scale farming to promote cultural heritage, and highly intense systems decrease and converge with low ones (e.g., organic, permaculture and regenerative farming) allowing for sustainable use of resources (Duarte et al., 2008). Even if organic farming may not always have higher biodiversity than comparable conventional farms (Dimambro et al., 2018; Tscharntke et al., 2021), an increase in biodiversity in organic systems is often attributed to the more heterogeneous landscapes (including crop diversity, boundary features and wooded areas). The beneficial impacts on the environment (Aldanondo-Ochoa and Almansa-Sáez, 2009; Gracia and de Magistris, 2008) are also related to cultivating lands without using mineral fertilisers and synthetic pesticides (IFOAM, 2023), that promotes pollination and biological pest control (Senapathi et al., 2015) in landscapes with a long tradition of extensive agriculture (Grass et al., 2019). Indeed, more emphasis is given to extensive grazing, meadows, hedgerows, small forest patches and forest hedges, that can support current culturally important agrobiodiversity, and improve connectivity. In Europe, these High Nature Value farmlands, defined in relation to land use, biodiversity and natural features (Andersen et al., 2003), also include silvopastoral agroforestry systems, and have been recognised for their high nature value (Parachinni et al., 2008), whose revitalization becomes a priority in NaC.

In the NfN narrative, the Renewable Energy Sources (RES) expansion is planned prioritising species and habitat conservation, avoiding areas important for biodiversity and maintaining the ecological integrity of landscapes (Fig. 3). On the other hand, in NaC the priority is given to the visual appeal of the landscapes, locating plants out of sight and far from culturally relevant places (Fig. 3). Among RES, biofuels energy is the most exploited solution in Europe (Bórawski et al., 2019). Indeed, they have the greatest relevance in NfS perspective (Fig. 3). However, according to the recent Paris Agreement Compatible Scenarios for Energy Infrastructures, demand for gaseous energy carriers will fall to less than a quarter of final energy demand in 2040, covered to just a minor extent by biofuels (CAN Europe, 2020). Moreover, producing biofuels from crops can harm the environment by decreasing biodiversity, polluting water, and it is not always associated with reducing $CO_2$ emissions (Cao & Pawlowski, 2013). Thus, tree plantation for biofuels is discouraged in the NfN perspective. Wind and solar energy are the others most adopted solutions (Bórawski et al., 2019). Although the achievement of 100% renewable energy scenario is feasible in Europe (Connolly et al., 2016; Potrc et al., 2021), the planning of RES expansion should take into account the impacts of the plants on nature and people and for this purpose different priorities have been pointed out in the NF narratives. In NfN, nature has the highest priority over RES expansion. It was documented that solar and wind plants can have direct and indirect impacts on species, and that land use impacts, caused by solar panels, may jeopardise their habitat (Gasparatos et al., 2017).Wind plants, which have less direct impacts on habitat, may cause the loss of their functional habitat due to species avoidance of disturbed areas (Gasparatos et al., 2017), and high collision risks for bats and birds (Gasparatos et al., 2017). Moreover, RES plants may have visual impacts, making the societal acceptance of

RES challenging (Gasparatos et al., 2017; Oudes & Stemke, 2021), which is taken into account in NaC view.

Urban sprawl, which refers to low-density expansion of large urban areas into the surrounding agricultural areas (EEA, 2004 ), is the main consequence of urbanisation (Sorensen, 1999; Habibi & Asadi, 2011; Yao et al., 2022). In NfN no increase in urban sprawl is expected and high-rise compact cities are developed, contrary to NaC perspective. This solution is proposed also worldwide to address the challenge of a growing population while limiting consumption of natural resources (Habibi & Asadi, 2011; Chen et al., 2016; Yao et al., 2022). In NfS sprawl is envisioned in peri-urban areas to improve contact between society and natural features and facilitate NCP provisioning. Indeed, peri-urban areas encompass different landscapes such as protected areas, forested hills, preserved woodlands, agricultural lands, and important wetlands that can provide ecosystem services to citizens (Douglas, 2006; Huang et al., 2009). Nevertheless it is important to enable a gradual and non aggressive urbanisation in these areas, within the constraint of the sustainable development imperative, in order to protect natural environment and NCP (Choy et al., 2008; Wandl & Magoni, 2017; Mortoja & Yigitcanlar, 2021). Moreover, the rural abandonment envisioned in NfN and NfS perspectives may lead to consequences in terms of conservation. However, this has increased conflicts between wildlife and people: the population growth of carnivore species has been associated with negative socio-economic impacts, such as economic loss due to livestock and crop damage, or negative emotions associated with the fear of attacks and damages to people and human activities (Gervasi et al., 2021; Rode et al., 2021).

In NfN green spaces are managed to protect ecological processes, improve connectivity to promote wildlife and plant dispersal (Mansur et al., 2022). Moreover, they promote urban rewilding, which is achieved by the implementation of soft management practices that lead to restore and improve the complexity of understory vegetation in city parks (Mansur et al., 2022). Restoration approaches can vary depending on the NF values. Passive restoration aligns more with NfN, emphasising natural regeneration. Active restoration is associated with NfS and NaC, since it involves human intervention (Atkinson & Bonsen, 2020). Rewilding is an European passive restoration strategy consisting in letting nature take care of itself, allowing natural processes to shape the land and sea, repair damaged ecosystems and restore degraded landscapes (Rewilding Europe, 2023) . Challenges for this approach in Europe include broad project scopes, monitoring difficulties, and the lack of evaluation frameworks, hindering rewilding's full potential. Overcoming these constraints is crucial for human and planetary health, benefiting biodiversity, food security, carbon sequestration, and restoring complex ecosystems (Svenning et al., 2016; Perino et al., 2019; Svenning, 2020).

In NaC, Green and Blue Infrastructures are mostly envisioned in urban areas, while in NfS they should be allocated across peri-urban areas and to connect cultivated lands, as they are associated with NCP, such as pollination (Liquete et al., 2015). In NfN, Green and Blue Infrastructure are important for rewilding marginal areas and enhancing connectivity in high-integrity landscapes. These elements, in addition to being associated with multiple benefits for humans, are also crucial in connecting protected areas (Liquete et al., 2015). As evidenced by the State of Nature Report, the Natura 2000 network was not efficient in maintaining habitat and species in a favourable condition, also because of disconnected sites across the network (EEA, 2020). Europe is planning to address this decline by building ‘a truly coherent Trans-European Nature Network’, which will be on the existing Natura 2000 network by analysing the potential connectivity between Natura 2000 sites using Green Infrastructure (EEA, 2020). In this light, the priorities provided by NF narratives are of paramount importance in finding consensus about Green Infrastructure that can support conservation and society development goals.

### References


Aldanondo-Ochoa, A. M., & Almansa-Sáez, C. (2009). The private provision of public environment: Consumer preferences for organic production systems. *Land Use Policy*, *26*(3), *669-682*. https://doi.org/10.1016/j.landusepol.2008.09.006

Andersen, E., Baldock, D., Bennett, H., Beaufoy, G., Bignal, E., Brouwer, F., ... & Zerva, G. (2003). Developing a high nature value indicator. *Report for the European Environment Agency, Copenhagen*. http://www.ieep.eu/assets/646/Developing_HNV_indicator.pdf

Armstrong, A. J., Holmes, C. M., & Henning, D. (2020). A changing world, again. How Appreciative Inquiry can guide our growth. Social Sciences & Humanities Open, 2(1), 100038. https://doi.org/10.1016/j.ssaho.2020.100038.

Atkinson, J., & Bonser, S. P. (2020). "Active" and "passive" ecological restoration strategies in meta-analysis. *Restoration Ecology, 28(5), 1032-1035*. https://doi.org/10.1111/rec.13229

Berbeć, A. K., Beata Feledyn-Szewczyk, B. (2018). Biodiversity of weeds and soil seed bank in organic and conventional farming systems. Institute of Soil Science and Plant Cultivation, State Research Institute in Puławy, Poland. https://doi.org/10.22616/rrd.24.2018.045

Boitani, L., & Linnell, J. D. (2015). Bringing large mammals back: large carnivores in Europe. *Rewilding European Landscapes, 67-84*. https://doi.org/10.1007/978-3-319-12039-3_4

Bórawski, P., Bełdycka-Bórawska, A., Szymańska, E. J., Jankowski, K. J., Dubis, B., & Dunn, J. W. (2019). Development of renewable energy sources market and biofuels in The European Union. Journal of cleaner production, 228, 467-484. https://doi.org/10.1016/j.jclepro.2019.04.242

Brown, J. (2010). The World Café: Shaping our future through conversations that matter. www.ReadHowYouWant.com accessed on 27th October 2023.

CAN Europe (2020). EEB technical summary of key elements. Building a Paris Agreement Compatible (PAC) energy scenario. https://www.pac-scenarios.eu/pac-scenario/scenario-development.html#Executivesummary

Cao, Y., & Pawłowski, L. (2013). Effect of biofuels on environment and sustainable development. *Ecological Chemistry and Engineering S, 20(4), 799-804.* https://doi.org/10.2478/eces-2013-0055

Chapron, G., Kaczensky, P., Linnell, J. D., Von Arx, M., Huber, D., Andrén, H., ... & Boitani, L. (2014). Recovery of large carnivores in Europe's modern human-dominated landscapes. *Science, 346(6216), 1517-1519.* https://doi.org/10.1126/science.1257553

Chaudhary, A., Pfister, S., & Hellweg, S. (2016). Spatially Explicit Analysis of Biodiversity Loss Due to Global Agriculture, Pasture and Forest Land Use from a Producer and Consumer Perspective. *Environmental Science & Technology 2016 50 (7), 3928-3936*. https://doi.org/10.1021/acs.est.5b06153

Chen, Y., Chen, Z., Xu, G., Tian, Z. (2016). Built-up land efficiency in urban China: Insights from the General Land Use Plan (2006–2020). *Habitat International, Vol. 51, Pag. 31-38.* https://doi.org/10.1016/j.habitatint.2015.10.014

Choy, D. L., & Sutherland, C. (2008). A changing peri-urban demographic landscape. *Australian Planner*, *45(3), 24-25.* https://doi.org/10.1080/07293682.2008.9982672

Cimatti, M., Ranc, N., Benítez-López, A., Maiorano, L., Boitani, L., Cagnacci, F., ... & Santini, L. (2021). Large carnivore expansion in Europe is associated with human population density and land cover changes. *Diversity and Distributions*, *27(4), 602-617.* https://doi.org/10.1111/ddi.13219

Clergue, B., Amiaud, B., & Plantureux, S. (2005). Assessment of biodiversity functions with agro-ecological indicators in agricultural areas. *Integrating Efficient Grassland Farming and Biodiversity*.

Connolly, D., Lund, H., & Mathiesen, B. V. (2016). Smart Energy Europe: The technical and economic impact of one potential 100% renewable energy scenario for the European Union. *Renewable and Sustainable Energy Reviews, 60, 1634-1653.* https://doi.org/10.1016/j.rser.2016.02.025

Dimambro, M., Rayns, F., Steiner, J., Carey, P. (2018). Countryside Stewardship organic management and conversion options: A scoping study to establish a monitoring protocol . Literature review. https://pure.coventry.ac.uk/ws/portalfiles/portal/40416921/Dimambro_Rayns_Steiner_and_Carey_2018.pdf

Douglas, I. (2006). Peri-Urban Ecosystem and Societies: Transitional Zones and Contrasting Values. In D. McGregor, D., Simon, & D. Thomson (Eds.). *The Peri-Urban Interface, pp. 18-27. London: Earthscan.* https://doi.org/10.4324/9781849775878

Duarte, F., Jones, N., Fleskens, L. (2008). Traditional olive orchards on sloping land: Sustainability or abandonment? *Journal of Environmental Management, Vol. 89, Issue 2, Pag. 86-98*. https://doi.org/10.1016/j.jenvman.2007.05.024

Dudley, N., & Alexander, S. (2017). Agriculture and biodiversity: a review. *Biodiversity, 18:2-3, 45-49.*https://doi.org/10.1080/14888386.2017.1351892

Dunn-Capper, R., Giergiczny, M., Fernández, N., Marder, F., & Pereira, H. M. (2024). Public preference for the rewilding framework: A choice experiment in the Oder Delta. People and Nature, pan3.10582. https://doi.org/10.1002/pan3.10582

EC (2016) No net land take by 2050? European Commission, Directorate-General for Environment, Publications Office of the European Union, 2016. https://data.europa.eu/doi/10.2779/537195.

EC (2020a). EU biodiversity strategy for 2030: bringing nature back into our lives. Communication from the commission to the european parliament, the council, the european economic and social committee and the committee of the regions. European Commission. COM 380. Brussels. https://data.europa.eu/doi/10.2779/048

EC (2022a). Proposal for a Regulation of the European Parliament and of the Council on Nature Restoration. Brussels, 22.6. 2022, COM 304 Final 2022/0195 (COD).

EC (2023b). European Climate Law. European Commission. https://climate.ec.europa.eu/eu-action/europeangreen-deal/european-climate-law_en accessed on 10th November 2023.

EEA (2004). Urban sprawl. EEA Glossary. https://www.eea.europa.eu/help/glossary/eea-glossary/urban-sprawl accessed on 19th November 2023.

EEA (2020). Building a coherent Trans-European Nature Network, 05/2020, European Environment Agency.

Finger, R., Swinton, S. M., El Benni, N., & Walter, A. (2019). Precision farming at the nexus of agricultural production and the environment. *Annual Review of Resource Economics*, *11*, *313-335*. https://doi.org/10.1146/annurev-resource-100518-093929

Fischer, J., Brosi, B., Daily, G.C., Ehrlich, P.R., Goldman, R., Goldstein, J., Lindenmayer, D.B., Manning, A.D., Mooney, H.A., Pejchar, L., Ranganathan, J. and Tallis, H. (2008). Should agricultural policies encourage land sparing or wildlife-friendly farming? *Frontiers in Ecology and the Environment, 6: 380-385.* https://doi.org/https://doi.org/10.1890/070019

Gasparatos, A., Doll, C. N., Esteban, M., Ahmed, A., & Olang, T. A. (2017). Renewable energy and biodiversity: Implications for transitioning to a Green Economy. *Renewable and Sustainable Energy Reviews*, *70*, *161-184*. https://doi.org/10.1016/j.rser.2016.08.030

Gervasi, V., Linnell, J. D., Berce, T., Boitani, L., Ciucci, P., Cretois, B., ... & Gimenez, O. (2021). Ecological correlates of large carnivore depredation on sheep in Europe. *Global Ecology and Conservation*, *30*, *e01798*. https://doi.org/10.1016/j.gecco.2021.e01798

Gracia, A., & De Magistris, T. (2008). The demand for organic foods in the South of Italy: A discrete choice model. *Food policy*, *33*(5), *386-396*. https://doi.org/10.1016/j.foodpol.2007.12.002

Grass, I., Loos, J., Baensch, S., … & Tscharntke, T. (2019). Land-sharing/-sparing connectivity landscapes for ecosystem services and biodiversity conservation. People Nat. 2019; 1: 262–272. https://doi.org/10.1002/pan3.21

Habibi, S., & Asadi, N. (2011). Causes, Results and Methods of Controlling Urban Sprawl. *Procedia Engineering, Vol. 21, Pag. 133-141.* https://doi.org/10.1016/j.proeng.2011.11.1996

Huang, S.-L., Wang, S.-H., Budd, W. B. (2009). Sprawl in Taipei's peri-urban zone: Responses to spatial planning and implications for adapting global environmental change, Landscape and Urban Planning, Vol. 90, Issues 1–2, Pag. 20-32, https://doi.org/10.1016/j.landurbplan.2008.10.010

IFOAM (2023). Principles of Organic Agriculture. https://www.ifoam.bio/principles-organic-agriculture-brochure accessed on 30th October 2023.

Liquete, C., Kleeschulte, S., Dige, G., Maes, J., Grizzetti, B., Olah, B., & Zulian, G. (2015). Mapping green infrastructure based on ecosystem services and ecological networks: A Pan-European case study. *Environmental Science & Policy, 54, 268-280.* https://doi.org/10.1016/j.envsci.2015.07.009

Mansur, A. V., McDonald, R. I., Güneralp, B., Kim, H., de Oliveira, J. A. P., Callaghan, C. T., ... & Pereira, H. M. (2022). Nature futures for the urban century: Integrating multiple values into urban management, *Environmental Science & Policy, Vol. 131, Pag. 46-56.* https://doi.org/10.1016/j.envsci.2022.01.013

Marcacci, G., Gremion, J., Mazenauer, J., Sori, T., Kebede, F., Ewnetu, M., ... & Jacot, A. (2020). Large-scale versus small-scale agriculture: Disentangling the relative effects of the farming system and semi-natural habitats on birds' habitat preferences in the Ethiopian highlands. *Agriculture, ecosystems & environment, 289, 106737.* https://doi.org/10.1016/j.agee.2019.106737

Mendelow, A. L., "Environmental Scanning--The Impact of the Stakeholder Concept" (1981). *ICIS 1981 Proceedings*. 20. https://aisel.aisnet.org/icis1981/20

Mortoja, M. G., & Yigitcanlar, T. (2022). Why is determining peri-urban area boundaries critical for sustainable urban development? *Journal of Environmental Planning and Management*, *66(1)*, *67-96*. https://doi.org/10.1080/09640568.2021.1978405

O'Connor L. M. J., Pollock L. J., Renaud J., Verhagen W., Verburg P. H. , Lavorel S., Maiorano L., Thuiller W. (2021). Balancing conservation priorities for nature and for people in Europe. Science. 372(6544):856-860. https://doi.org/10.1126/science.abc4896

O'Farrell, P. J., & Anderson, P. M. (2010). Sustainable multifunctional landscapes: a review to implementation. *Current Opinion in Environmental Sustainability*, *2*(1-2), 59-65. https://doi.org/10.1016/j.cosust.2010.02.005

Oudes, D., & Stremke, S. (2021). Next generation solar power plants? A comparative analysis of frontrunner solar landscapes in Europe. *Renewable and Sustainable Energy Reviews, 145, 111101*. https://doi.org/10.1016/j.rser.2021.111101

Paracchini, M., Petersen, J., Hoogeveen, Y., Bamps, C., Burfield, I., Van Swaay, C. (2008). High Nature Value Farmland in Europe - An Estimate of the Distribution Patterns on the Basis of Land Cover and Biodiversity Data. *Publications Office, Luxembourg.* https://data.europa.eu/doi/10.2788/8891

Perino, A., Pereira, H. M., Navarro, L. M., Fernández, N., Bullock, J. M., Ceaușu, S.,... & Wheeler, H. C. (2019). Rewilding complex ecosystems. *Science*, *364(6438)*, *eaav5570*. https://doi.org/10.1126/science.aav5570

Pingali, P. L. (2012). Green revolution: impacts, limits, and the path ahead. *Proceedings of the national academy of sciences*, *109*(31*), 12302-12308.* https://doi.org/10.1073/pnas.0912953109

Potrč, S., Čuček, L., Martin, M., & Kravanja, Z. (2021). Sustainable renewable energy supply networks optimization–The gradual transition to a renewable energy system within the European Union by 2050. *Renewable and Sustainable Energy Reviews, 146, 111186.* https://doi.org/10.1016/j.rser.2021.111186

Raj, E. F. I., Appadurai, M., Athiappan, K. (2021). Precision Farming in Modern Agriculture. In: Choudhury, A., Biswas, A., Singh, T.P., Ghosh, S.K. (eds). Smart Agriculture Automation Using Advanced Technologies. Transactions on Computer Systems and Networks. *Springer, Singapore*. https://doi.org/10.1007/978-981-16-6124-2_4

Rewilding Europe (2023). www.rewildingeurope.com accessed on 18th November 2023.

Rode, J., Flinzberger, L., Karutz, R., Berghoefer, A., & Schroeter-Schlaack, C. (2021). Why so negative? Exploring the socio-economic impacts of large carnivores from a European perspective. *Biological Conservation, 255, 108918*. https://doi.org/10.1016/j.biocon.2020.108918

Senapathi, D., Jacobus, B. C., Breeze, T. D., David, K., Potts, S. G., and Carvalheiro, L. G. (2015). Pollinator conservation: the difference between managing for pollination services and preserving pollinator diversity. *Current Opinion in Insect Science, 12. pp. 93-101*. https://doi.org/10.1016/j.cois.2015.11.002

Svenning, J. C., Pedersen, P. B., Donlan, C. J., Ejrnæs, R., Faurby, S., Galetti, M., Hansen, D. M., Sandel, B., Sandom, C. J., Terborgh, J. W., Vera, F. W. (2016). Science for a wilder Anthropocene: Synthesis and future directions for trophic rewilding research. *Proc Natl Acad Sci U S A;113(4) :898-906*. https://doi.org/10.1073/pnas.1502556112

Svenning, J-C. (2020). Rewilding should be central to global restoration efforts, *One Earth, Volume 3, Issue 6, Pages 657-660*. https://doi.org/10.101/j.oneear.2020.11.014

Troiano, S., Carzedda, M. & Marangon, F. (2023). Better richer than environmentally friendly? Describing preferences toward and factors affecting precision agriculture adoption in Italy. *Agric Econ 11, 16*. https://doi.org/10.1186/s40100-023-00247-w

Tscharntke, T., Grass, I., Wanger, T. C., Westphal, C., Batáry, P. (2021). Beyond organic farming – harnessing biodiversity-friendly landscapes. *Trends in Ecology & Evolution, Vol. 36, Issue 10, Pag. 919-930*. https://doi.org/10.1016/j.tree.2021.06.010

Yao, Y., Haozhi, P., Cui, X., Wang, Z. (2022). Do compact cities have higher efficiencies of agglomeration economies? A dynamic panel model with compactness indicators. *Land Use Policy, Vol. 115, 106005*. https://doi.org/10.1016/j.landusepol.2022.106005

Wandl, A., & Magoni, M. (2017). Sustainable Planning of Peri-Urban Areas: Introduction to the Special Issue, *Planning Practice & Research, 32:1, 1-3.* https://doi.org/10.1080/02697459.2017.1264191
